# Spatiotemporal characterization of glial cell activation in an Alzheimer’s disease model by spatially resolved transcriptome

**DOI:** 10.1101/2021.06.28.450154

**Authors:** Hongyoon Choi, Eun Ji Lee, Jin Seop Shin, Hyun Kim, Sungwoo Bae, Yoori Choi, Dong Soo Lee

**Affiliations:** Department of Nuclear Medicine, Seoul National University College of Medicine, Republic of Korea; Department of Nuclear Medicine, Seoul National University Hospital, Seoul, Republic of Korea; Department of Molecular Medicine and Biopharmaceutical Sciences, Graduate School of Convergence Science and Technology, Seoul National University, Seoul, Republic of Korea

**Author notes:** [Correspondence and Reprint Request] Dong Soo Lee, MD.,Ph.D., Department of Nuclear Medicine, Seoul National University College of Medicine 101 Daehak-ro, Jongno-gu, Seoul 03080, Korea, Yoori Choi, Ph.D., Department of Nuclear Medicine, Seoul National University College of Medicine 101 Daehak-ro, Jongno-gu, Seoul 03080, Korea, Hongyoon Choi, MD., Ph.D., Department of Nuclear Medicine, Seoul National University Hospital 101 Daehak-ro, Jongno-gu, Seoul 03080, Korea. These authors contributed equally to this work (Hongyoon Choi, Eun Ji Lee).

**Keywords:** Alzheimer’s disease, Spatially resolved transcriptome, Amyloid plaques, Microglia, Astrocytes, reactive glial cells

## Abstract

The pathophysiological changes that occur with the progression of Alzheimer’s disease (AD) are well known, but understanding the spatiotemporal heterogeneity of the brain is needed. Here, we investigated the spatially resolved transcriptome in a 5XFAD AD model of different ages to understand regional changes at the molecular level. We identified early alterations in the white matter (WM) of the AD model before the definite accumulation of amyloid plaques in the gray matter (GM). Changes in the early stage of the disease were involved primarily in glial cell activation in WM, whereas the changes were prominent in the later stage of pathology in GM. We confirmed that disease-associated microglia (DAM) and astrocyte (DAA) signatures also showed initial changes in WM and that activation spreads to GM. Trajectory inference using microglial gene sets revealed the subdivision of DAMs with different spatial patterns. Taken together, these results help to understand the spatiotemporal changes associated with reactive glial cells as a major pathophysiology of AD and provide information for diagnosis and prognosis based on spatiotemporal changes caused by amyloid accumulation in AD.

## INTRODUCTION

Advances in single-cell analysis have revealed the diversity of brain cells, and the dynamics of cellular changes in Alzheimer’s disease (AD) have been observed^1–3^. Single-nucleus transcriptome analysis of patients with AD showed major changes characterized by myelination, inflammation, and neuronal survival^2^. In addition, single-cell analysis of AD models especially observed molecular changes in microglia and astrocytes. Microglia respond to β-amyloid (Aβ) plaques, and genetic changes in activated response microglia (ARM) are associated with AD risk genes^4^. Based on these genetic changes in microglia that are specific to AD, disease-associated microglia (DAM) are defined as a microglial subtype^5,6^. The alteration of specific subtypes of astrocytes in AD was also identified and defined as disease-associated astrocytes (DAAs) ^7^. Despite the identification of the cellular landscape according to AD pathophysiology in specific brain regions, the limitation is the loss of spatial information on cellular networks. Because of the diversity of cellular profiles according to brain regions^8–10^, it is unclear in which brain regions the cellular changes identified in AD are distributed and how they change across brain regions as the disease progresses.

Molecular changes in brain cells as AD pathology progresses should consider regional heterogeneity. According to recent studies, microglia are composed of various subtypes and show high plasticity depending on the surrounding environment^8,11,12^. In the AD model, white matter-associated microglia (WAM), a type of microglia specific only to the white matter (WM), play an important role in the clearance of myelin but do not exist near amyloid plaques^13^. However, the other subtypes of microglia are present and play different roles in gray matter (GM) ^13^. In addition, macrophages exist in different subtypes depending on their location, such as the dura mater, subdural meninges, and choroid plexus^14^, and differential patterns of astrocytes and neurons are observed according to the cortical layer^15,16^. These studies show that not only differences in cell subtypes but also functional differences according to brain regions such as GM and WM can be observed. Since brain cells have diversity and dynamics in inter- and intraregions, spatial information within neural circuits is important, and understanding the pathophysiology of AD requires spatiotemporal landscape studies at the molecular level throughout the brain.

Here, we applied spatially resolved transcriptomic analysis in 5XFAD AD models of different ages to identify spatiotemporal patterns of disease progression. First, distinctive brain regions clustered by gene expression were identified, and then molecular changes were analyzed in early (3-month-old) and late (7-month-old) stage AD models. As a result, the analysis confirmed gene patterns that change according to disease progression in each brain region and initial molecular changes related to glial cell activation in WM before the changes in GM. In addition, the spatiotemporal trajectories of the microglia-related gene signature in the spot revealed distinctive activation patterns and found each major marker gene set. These results provide spatiotemporal molecular profiles in the pathophysiology of AD and distinctive activation patterns of microglia and astrocytes that change with AD progression.

## RESULTS

### Distinctive gene expression patterns commonly found in the GM of AD by spatial transcriptome-based cluster analysis

Spatial transcriptomic data of the AD model of 5XFAD and age-matched wild-type (WT) mice were obtained at 3 and 7 months of age. Notably, a 7.5-month-old AD model showed amyloid plaque in GM, while the 2-month-old AD model did not. The 4-month-old AD model showed a small number of amyloid plaques in the thalamus and cortex (**Supplementary Fig. 1a**). In particular, as a result of confirming the amyloid deposition in the brain tissues of the AD model, which were the same as those for the spatial transcriptomic data, a small proportion of amyloid deposition was observed in the 3-month-old AD model, and amyloid plaques were also observed in the 7-month-old AD model (**Supplementary Fig. 1b**). A total of 12,247 spots containing 32,285 mRNA expression data points were clustered according to the expression patterns only (detailed methods are described in Materials and Methods). Accordingly, 15 different clusters were identified. These clusters corresponded to anatomical structures (**Figure 1a**). For example, cluster 6 represented the cerebral cortex including mainly outer layers, and cluster 3 represented the cerebral cortex including mainly inner layers. Cluster 4 included the hippocampus, and cluster 9 represented the striatum (**Supplementary Table 1**). The expression data of spots were represented by Uniform Manifold Approximation and Projection (UMAP) plots, a dimension reduction method for visualization17 (**Figure 1b**). Spots with different mice are also depicted (**Figure 1c**). Markers of each cluster were extracted and visualized with a heatmap (**Supplementary Fig. 2**).

**Figure 1.**
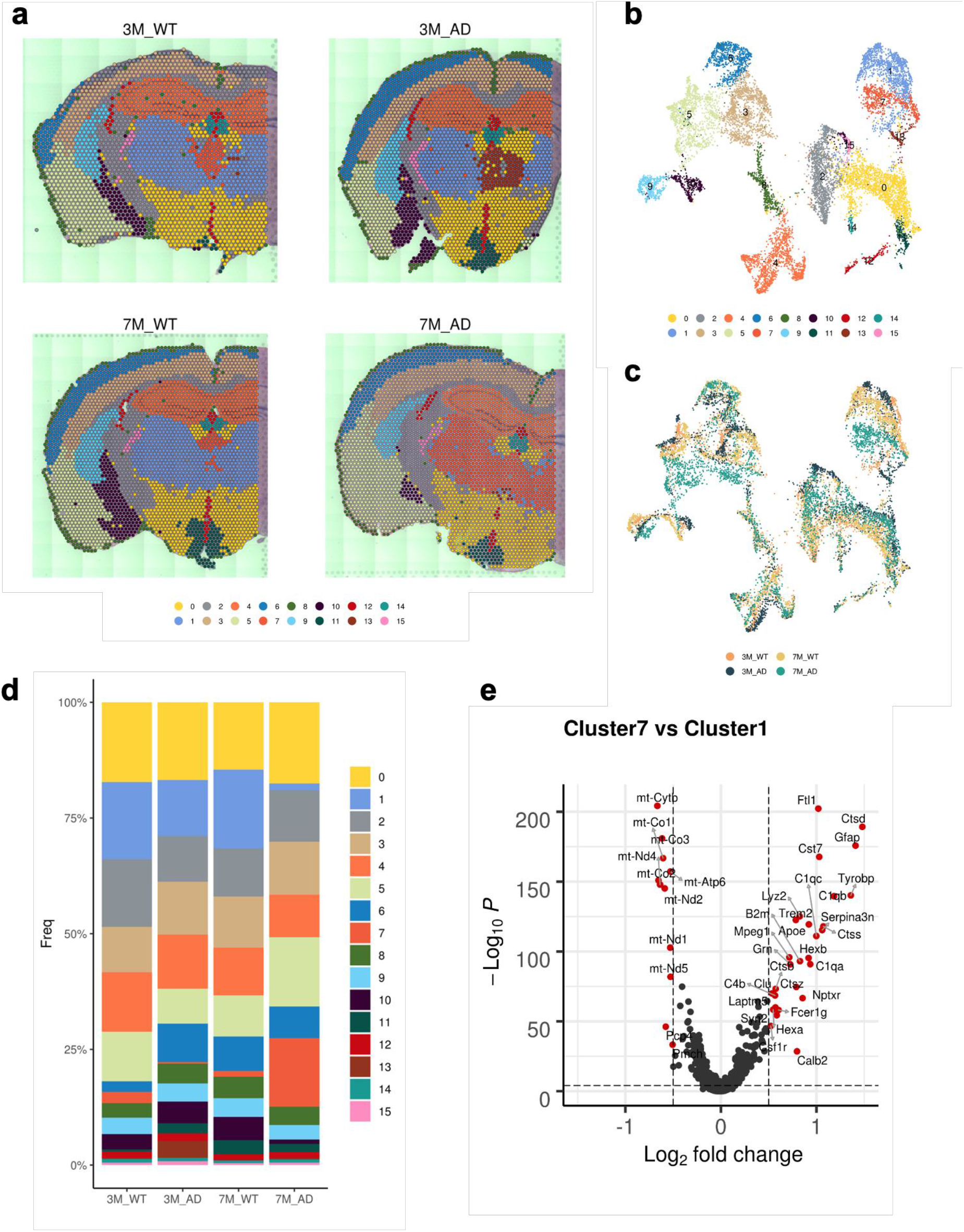
Clusters of spots of spatially resolved transcriptomes in WT and AD models. (a) Fifteen clusters were identified according to transcriptomic data of 3- and 7-month-old WT and AD models. (b) A two-dimensional reduction map, UMAP, colored with clusters is depicted. Each dot represents transcriptomic data of each spot. (c) A UMAP colored with mouse types was depicted. (d) The frequency of spots of each cluster for different mice is represented. Notably, cluster 7 was especially more frequent in the 7-month-old AD model. (e) As clusters 7 and 1 represented the thalamus while cluster 7 was dominantly found in the 7-month-old AD model, differentially expressed genes between the two clusters were identified. A volcano plot represents these differentially expressed genes. A positive log-fold change represented upregulated genes in cluster 7. (WT: wild type; AD: Alzheimer’s disease; 3M: 3-month-old; 7M: 7-month-old)

The frequency of each cluster was represented to identify a specific cluster enriched in the AD models (**Figure 1d**). Cluster 7 was markedly increased in the thalamus of AD model at 7 months. However, cluster 1, which represented the thalamus in other mice, was decreased in the AD model at 7 months. This finding was also identified in **Figure 1a**, which shows a change in the thalamus to the gene expression pattern from cluster 1 to 7 in the AD model at 7 months. As clusters 1 and 7 represented the thalamus but were differentially found in the 7-month-old AD model, differentially expressed genes (DEGs) between these two clusters were analyzed (**Figure 1e**). The genes highly expressed in cluster 7 compared with cluster 1 included *Ctsd, Gfap, Tyrobp, C1qb, Ctss, Serpina3n, Cst7, Ftl1, C1qc, C1qa, Trem2,* and *Hexb.* These genes were associated with myeloid leukocyte activation, microglial activation, glial cell activation, and lysosome pathways according to gene ontology (GO) and Kyoto Encyclopedia of Genes and Genomes (KEGG) pathway analyses (**Supplementary Fig. 3a**). Relatively decreased expression in cluster 7 included mitochondrial genes such as *mt-Cytb* and *mt-Co1*. These genes were associated with ribonucleotide metabolic processes and ATP metabolic processes according to GO analysis (**Supplementary Fig. 3b**). Immunofluorescence (IF) images of the thalamus showed increased Iba1, GFAP, and Lamp1 expression, representing glial cell accumulation and lysosomal activity, in the thalamus of the AD model after 4 months (**Supplementary Fig. 3c**).

We then analyzed DEGs in specific brain regions of the AD brain. First, genes differentially expressed in the cerebral cortex of AD, cluster 3, were identified (**Figure 2a**). UMAP showed spots according to the origin of the samples (**Figure 2b**). In cluster 3, differential gene expression was found mainly in the AD model at 7 months, while mice at 3 months showed few differentially expressed genes (**Figure 2c, Supplementary Fig. 4a**). The genes highly expressed in the 7-month-old AD model included *Gfap, Ftl1, Tyrobp, Ctsd, Cst7, Hexb,* and *C1qb*, which were related to inflammation mediated by microglia and astrocytes (**Figure 2c**). These upregulated genes were similar to the markers related to cluster 7, included mainly in the thalamus of the 7-month-old AD model (**Figure 2d**). GO pathway analysis revealed that upregulated genes in the cerebral cortex of the AD model at 7 months were associated with gliogenesis and neuroinflammation related to microglia and astrocytes (**Figure 2e**). In addition, lysosome, apoptosis, and phagosome activity was enriched in the cerebral cortex of 7-month-old AD model. Genes downregulated in the AD model compared with WT included *Pmch, Arf5, BC004004, Trbc2,* and *Arc.* These genes were associated with Parkinson’s disease, amyotrophic lateral sclerosis, Alzheimer’s disease and oxidative phosphorylation in KEGG pathways (**Supplementary Fig. 4b**). Indeed, IF images showed increased activity of microglia, astrocytes, and lysosomal function in the cerebral cortex using antibodies against Iba1, GFAP, and Lamp1 from AD model older than 4 months (**Supplementary Fig. 4c**). We additionally tested DEGs in the AD brain according to the regions. Accordingly, the hippocampus (cluster 4), striatum (cluster 9), and outer cortex (cluster 6) shared genes differentially expressed at 7 months (**Figure 2f**, **Supplementary Fig. 5**). The commonly upregulated genes in these GM regions of the 7-month-old AD model are represented in **Figure 2f**.

**Figure 2.**
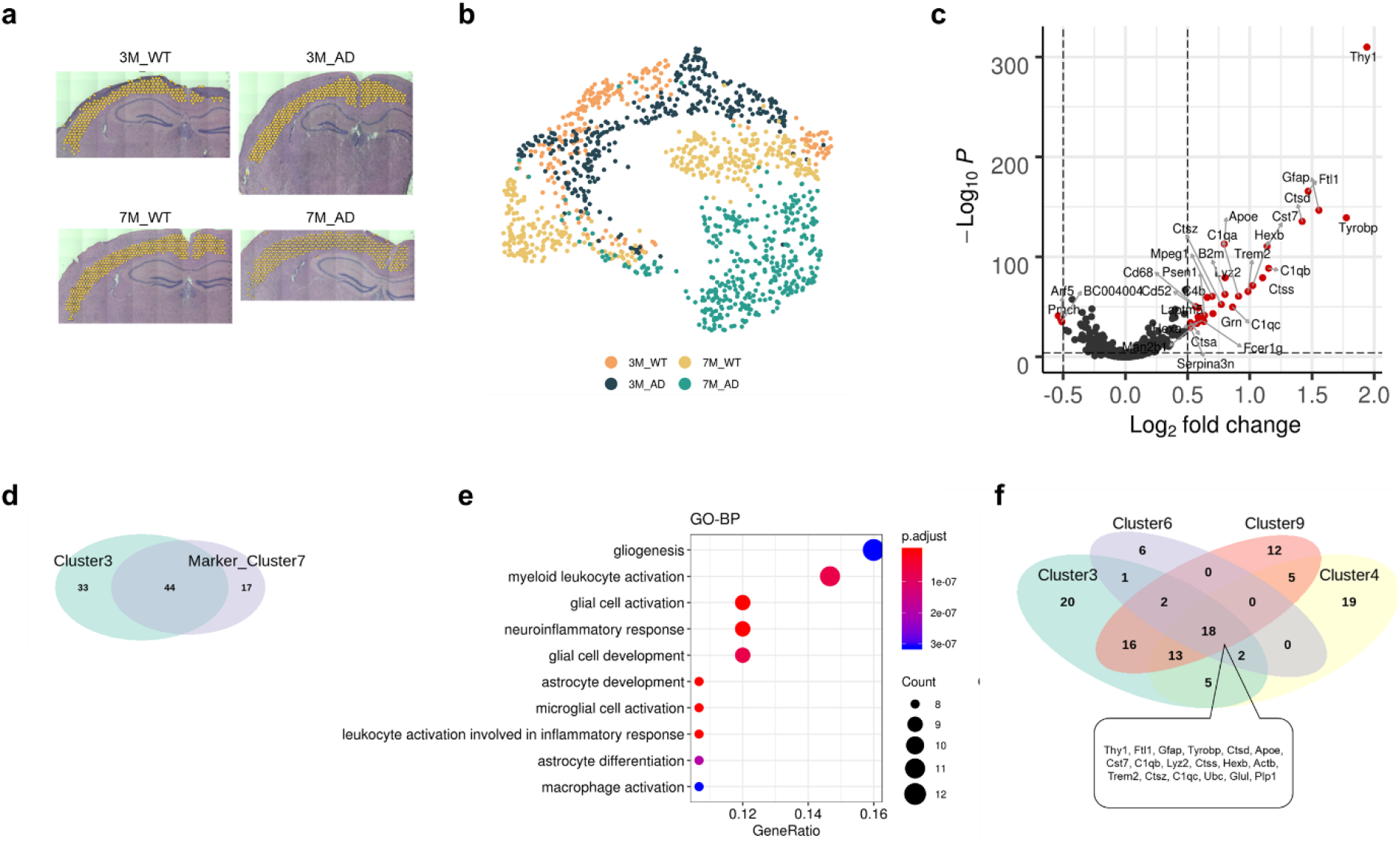
Genes differentially expressed in the gray matter of the AD model. (a) As a cluster of the cerebral cortex, cluster 3 was selected to find differentially expressed genes in AD compared with WT. (b) UMAP represented spots of cluster 3 according to the origin of samples. (c) A volcano plot depicting differentially expressed genes of cluster 3 in the AD model compared with the WT at 7 months. (d) The upregulated genes in the 7-month-old AD model were similar to the markers of cluster 7, which occupied the thalamus of the 7-month-old AD model. (e) GO terms related to upregulated genes in cluster 3 of 7-month-old AD models are represented. (f) A Venn diagram representing the upregulated genes in the 7-month-old AD model for different clusters (clusters 3, 4, 6, and 9). There were 18 common upregulated genes in all these GM clusters.

### Distinctive spatial pattern of DEGs within WM in early AD

Cluster 2, which included mainly WM, showed distinctive patterns in DEGs in AD (**Figure 3a**). A UMAP represented spots of cluster 2 (**Figure 3b**). In cluster 2, striking DEGs were identified in 3-month-old mice as well as 7-month-old mice (**Figure 3c**). At 3 months, the AD model showed that many genes were upregulated compared with the WT. These genes included *Tmem242, Ctss, Hsd17b12, Mrpl51, Tial1, Gm10076, Cops4, Bloc1s1, Nr2f2, Llph, Mag,* and *Arpc1b.* Additionally, many genes were downregulated compared with WT. These genes included *Camkk1, Snd1, Metap1d, Itpa, Rab15, Bc1, Cyfip1, Fam3c, Nrd1,* and *Trak1*. The upregulated genes in the WM of 3-month-old AD brains were associated with neuron projection development, gliogenesis, and ensheathment of neurons (**Figure 3d**). qPCR analysis also showed that the mRNAs encoding *Mag* and *MOG* were significantly increased in the 3-month-old AD model compared to the control and 7-month-old AD model (**Supplementary Fig. 6a**). KEGG pathways showed increased enrichment in ribosome, endocytosis, and MAPK signaling pathways. However, the decreased genes in the WM of the 3-month-old AD model were associated with synapse organization, regulation of membrane potential, and neuron death (**Figure 3e**). IF images with GFAP showed increased reactive astrocytes in the WM, internal capsule and corpus callosum of 4-month-old AD model. The lysosomal function shown by staining with anti-Lamp1 also showed the same pattern of change. Reactive microglia were gradually increased in the WM, while they were increased in 7.5-month-old AD model, and a few fluorescence activities were identified in the 4-month-old AD model (**Supplementary Fig. 6b**). At 7 months, many DEGs were also identified in cluster 2. Increased expression was found in *Ftl1, Ctsd, Apoe, C4b, Tyrobp,* and *Trem2*, which were similar to upregulated genes in other clusters of GM, such as the cerebral cortex, thalamus, hippocampus and striatum. qPCR analysis revealed significant increases in *Cst3, Trem2, and Tyrobp* in both WM and GM at 7 months in the AD model, and the upregulated expression of *Apoe* was identified only in WM (**Supplementary Fig. 6a**). At 7 months, enriched GO terms included microglial cell activation as well as ensheathment of neurons and myelination (**Supplementary Fig. 7**). Accordingly, genes differentially expressed in the WM of 3-month-old AD model were distinctive from others, particularly in the 7-month-old AD model identified in GM (cluster 3). However, genes differentially expressed in the WM of 7-month-old AD model included the common genes differentially expressed in the GM (**Figure 3f**). DEGs of each cluster are summarized in **Supplementary Table 2**.

**Figure 3.**
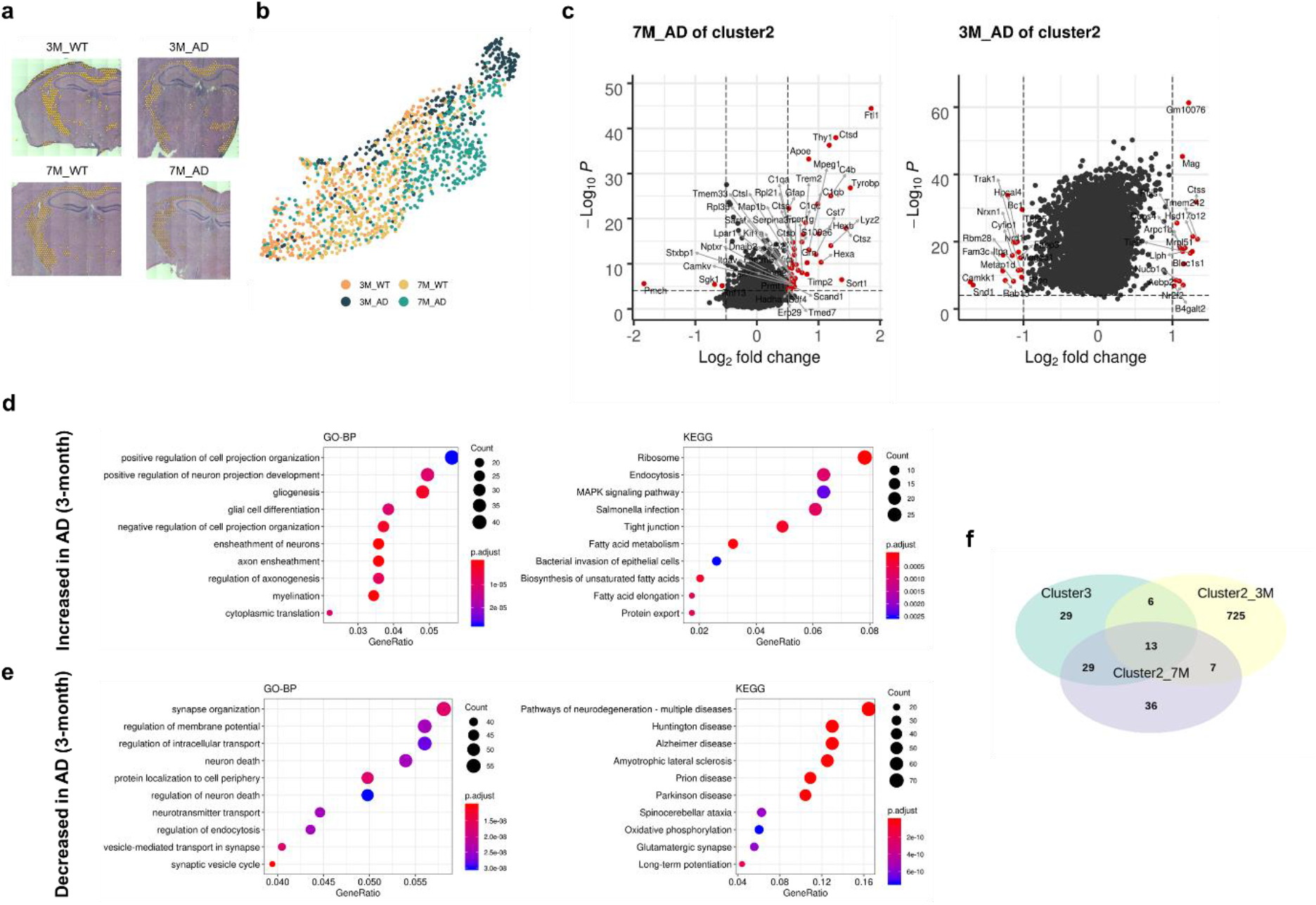
Genes differentially expressed in the WM of the AD model as an early change. (a) Cluster 2 represents WM regardless of mouse type. (b) UMAP represents transcriptomic profiles of cluster 2 according to mouse type. (c) Many differentially expressed genes were identified in the AD model at 3 months and at 7 months. (d) As an early change in WM (i.e., 3-month-old AD), upregulated genes were related to neuronal projection, gliogenesis, and axon ensheathment. (e) Downregulated genes in the 3-month-old AD model were related to synapse organization and membrane potential. (f) The upregulated genes of cluster 2 in the 3-month-old AD were different from the upregulated genes of GM in the 7-month-old AD. However, the upregulated genes of cluster 2 in the 7-month-old AD shared those of GM.

### Integrative analysis with the immunofluorescence image of Aβ

As the 7-month-old AD model showed amyloid plaques on the IF image, we integrated this image with spatial transcriptomic data. The image was spatially registered with the H&E image of the AD brain at 7 months using nonlinear transformation (**Supplementary Fig. 8a**). Therefore, we obtained registered images of amyloid plaques that corresponded to spots of spatial transcriptomic data using adjacent slides of the same brain tissue used for sequencing. Molecular features spatially correlated with the amyloid plaque image patterns were estimated using the spatial gene expression patterns by deep learning of tissue images (SPADE) tool18. Briefly, the image patch corresponding to the spot was extracted to estimate image features derived by a convolutional neural network model, and then correlated genes were identified. The first principal component (‘ImageLatent_1’) of the CNN-derived image features was mapped (**Figure 4a**). The spatial distribution of ‘ImageLatent_1’ corresponded visually to the pattern of amyloid deposits, while other principal components were different from the degree of amyloid deposits (**Supplementary Fig. 8b**). The top 12 genes spatially correlated with ‘ImageLatent_1’ are represented (**Figure 4b**). According to GO analysis, these genes represented the regulation of ion transport and myeloid leukocyte activation (Figure 4c). Genes associated with amyloid plaque image patterns (SPADE genes) partly overlapped with DEGs at 7 months in the cortex (**Figure 4d**). These genes, *Thy1, Gfap, Tyrobp, Ctsd, Cst7, C1qb, Lyz2, Ctss, Hexb, Trem2, B2m, C1qa, Mpreg1, Ctsz, C1qc, Cd68, Grn, Laptm5, Hexa,* and *Serpina3n*, were increased in 7-month-old AD model in the cortex and associated with amyloid plaque image patterns.

**Figure 4.**
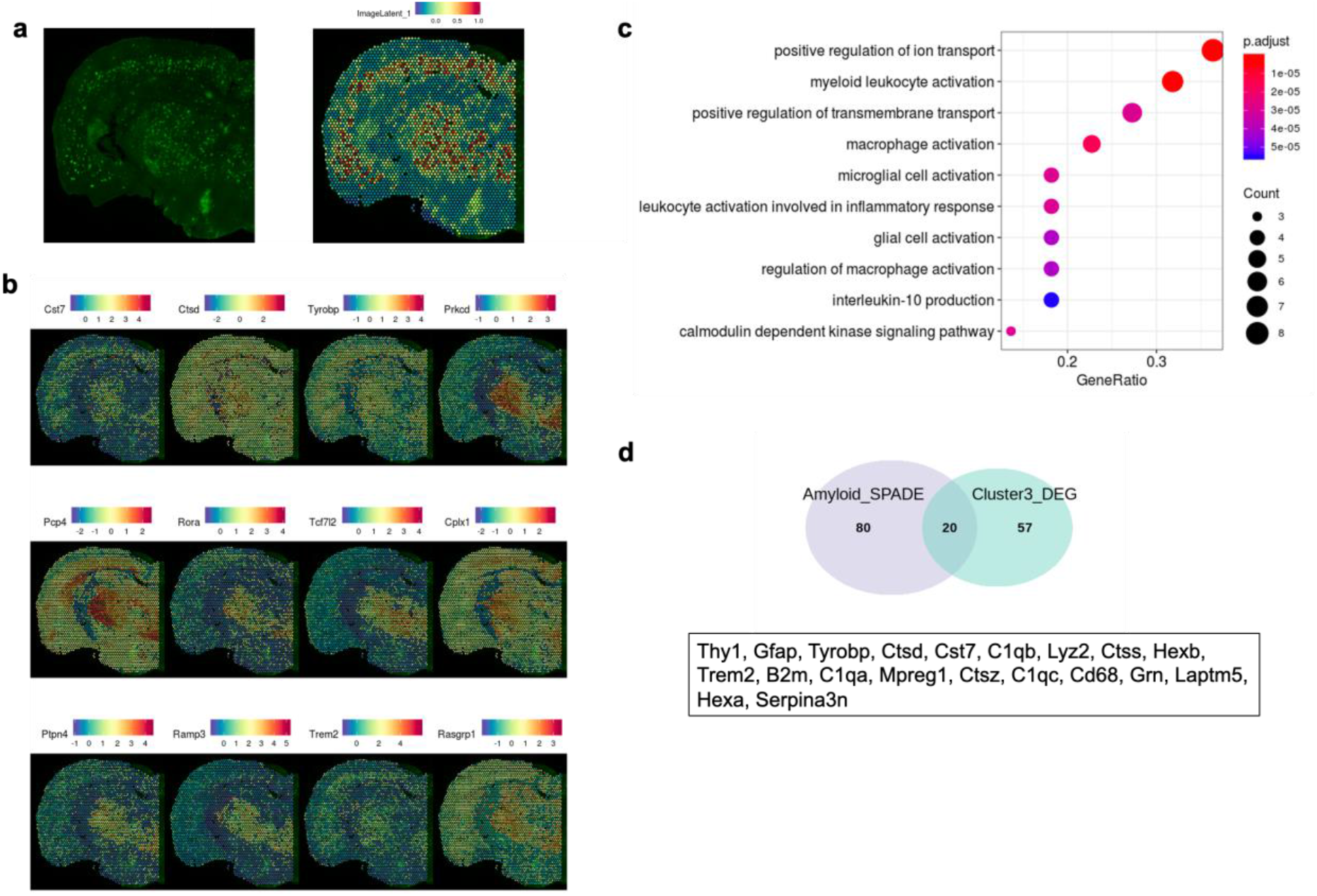
SPADE analysis showed genes spatially associated with amyloid plaque images. (a) Anti-amyloid plaque IF image in a 7-month-old AD model used for sequencing was spatially registered to the histology image of the spatially resolved transcriptome. The latent image (‘ImageLatent1’) derived by the first principal component of deep learning-based features is represented (right). (b) Top 12 genes spatially associated with ‘ImageLatent1’, analyzed by SPADE analysis. (c) GO terms of the spatially associated genes are represented. (d) Among genes spatially associated with ‘ImageLatent1’, 20 genes were upregulated genes in cluster 3 of 7-month-old AD model. The spatial distribution of these genes is presented.

### Spatial distribution of DAM and DAA signatures

Spatial transcriptomic data revealed that AD-related transcriptomic changes at 3 months involved mainly the WM, and then the features of reactive glial cells commonly extended to the cortex, thalamus, striatum and hippocampus. Thus, we analyzed disease-associated activation signatures of microglia and astrocytes, major players in AD-related neuroinflammation, in terms of spatial and temporal patterns. Spatial patterns of activated microglia, represented by the DAM signature, were identified. A module score of the DAM signature (DAMscore) was calculated by using the expression values of *Lpl, Cst7, Axl, Itgax, Spp1, Cd9, Ccl6*, and *Csf1* for each spot^5,6^ (**Supplementary Fig. 9a)**. The qPCR results showed significantly increased *Axl, Itgax,* and *Cst7* levels in both WM and GM in the 7-month-old AD model (**Supplementary Fig. 9b**). DAM scores of spots were represented by UMAP plots, which revealed spots with increased DAM scores in the 7-month-old AD model (**Figure 5a**). DAM scores of clusters were compared according to the mouse types (**Figure 5b**). In addition, the spatial distribution of DAM scores was represented (**Figure 5c**). Notably, these results showed that the DAM score was high in the WM of 3-month-old AD model (i.e., cluster 2). IF images with anti-Iba1 revealed that microglia in the internal capsule and corpus callosum were increased at 4 months in the AD model, and they were clearly increased in the cortex and thalamus at 7.5 months (**Supplementary Fig. 9c**). In addition, WT mice also showed relatively high DAM scores in brain regions such as the subdural area (cluster 8) and periventricular area (cluster 12) (**Figure 5c, Supplementary Fig. 10**), which suggested the heterogeneity of myeloid cells in the brain, identified as border-associated macrophages (BAMs) that substantially share gene sets of DAM^14^.

**Figure 5.**
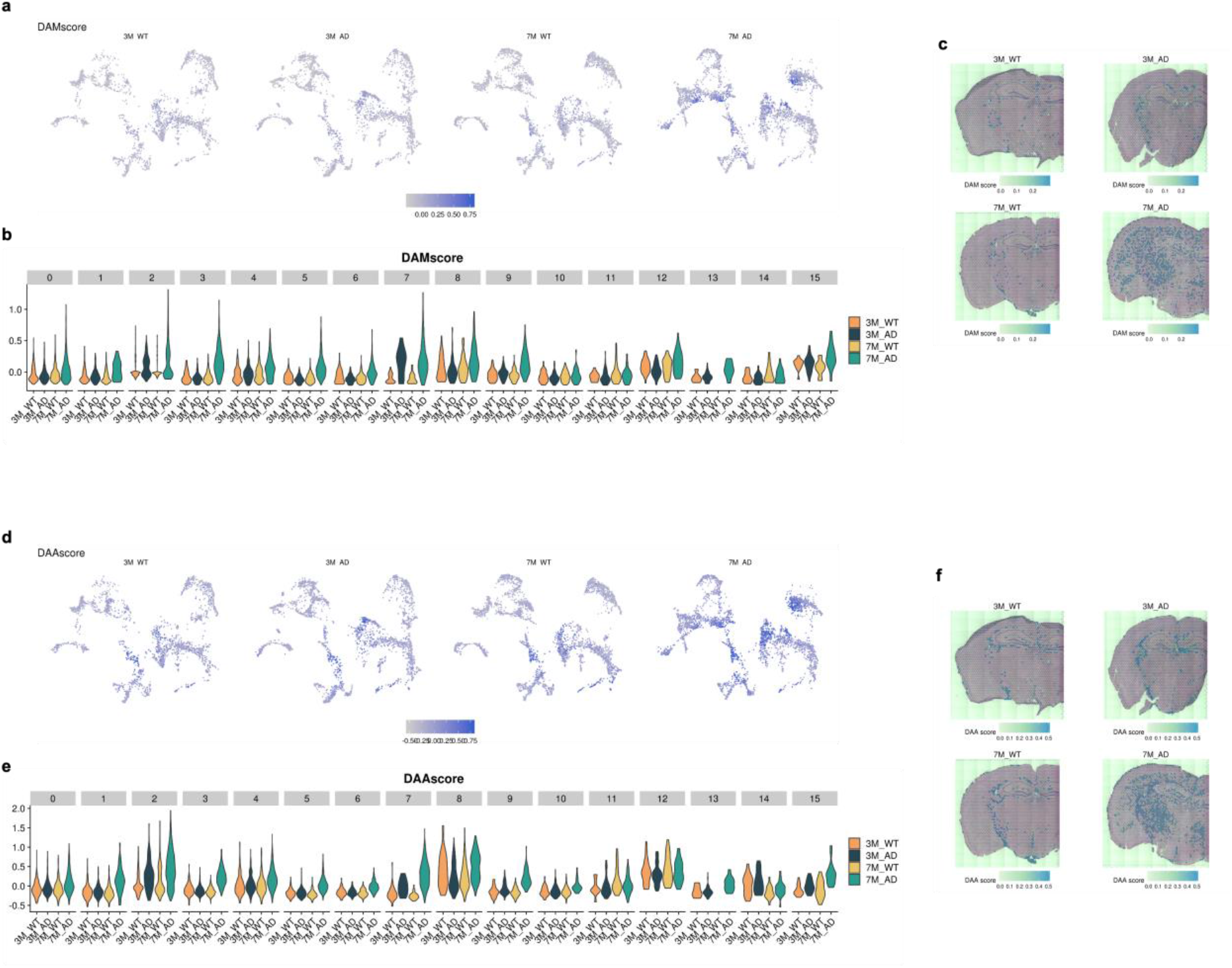
Spatial distribution of DAM and DAA signatures. (a) DAM score was represented by UMAP plots. (b) DAM scores of each cluster were compared and overall increased in 7-month-old AD model. The DAM score was also increased in the 3-month-old AD model in cluster 2. (c) The spatial distribution of DAM score showed an increase in the GM of 7-month-old AD and a relatively prominent increase in the WM of 3-month-old AD. (d) DAA score was represented by UMAP plots. (e) DAA score of each cluster were compared and overall increased in 7-month-old AD, dramatically in cluster 2. The DAA score was also increased in the 3-month-old AD model in cluster 2. (f) The spatial distribution of DAA score showed an increase in the 7-month-old AD model, particularly in the WM and thalamus regions and the slight increase in WM region in 3-month-old AD model.

We also analyzed the age-dependent spatial pattern of astrocytes activated only in AD, indicated by DAA signatures. The module score of the DAA signature (DAA score) was expressed in patterns of *Ggta1*, *Gsn*, *Osmr*, *Vim*, *Serpina3n*, *Ctsb*, and *Gfap*^7^. The DAA score increased at 3 months in the WM (i.e., cluster 2), and further increases were identified at 7 months in areas such as the thalamus and cortical cortex (i.e., clusters 1 and 3) (**Figure 5d, e**). Increased reactivity of astrocytes was observed, especially in the cortical cortex and thalamus at 7 months (**Figure 5f**). qPCR analysis revealed that *Gfap* and *Serpina3n* were significantly upregulated in both WM and GM in the 7-month-old AD model, while *Vim* showed a tendency of increased expression levels only in WM (**Supplementary Fig. 11a**). IF images of GFAP showed increased expression levels in the internal capsule and corpus callosum of a 4-month-old AD model. At 7 months, in addition to WM, Gfap expression was increased in the thalamus and cortex (**Supplementary Fig. 11b**). Thus, similar changes in spatial patterns were identified for the pathological conditions of DAM and DAA.

### Spatiotemporal reactive microglial patterns identified by trajectory analysis

Genes related to reactive microglia and homeostatic microglia were selected to find spatiotemporal patterns of microglial signatures in the brain (the gene sets are summarized in **Supplementary Table 3**) ^5,6^. Using these gene sets, trajectory of spots was inferred using Monocle 3^19^. UMAP plots based on the microglial gene sets depicted heterogeneous distribution in terms of the clusters representing anatomical regions (**Figure 6a**). In addition, spots of 7-month-old AD showed a trend of clustering in the center of the UMAP plot compared with the spots of other brains (**Figure 6b**). Accordingly, the direction of the trajectory of microglial activation according to disease progression could be inferred. The expression of key genes in microglia is presented in UMAP plots (**Supplementary Fig. 12**). These plots revealed that the expression of activated microglial genes, including *Trem2*, *Cst7*, and *Ccl6*, was increased according to the trajectory. According to the distribution of activated microglial genes and brain samples, four different trajectories were defined based on trajectory analysis (**Figure 6c**).

**Figure 6.**
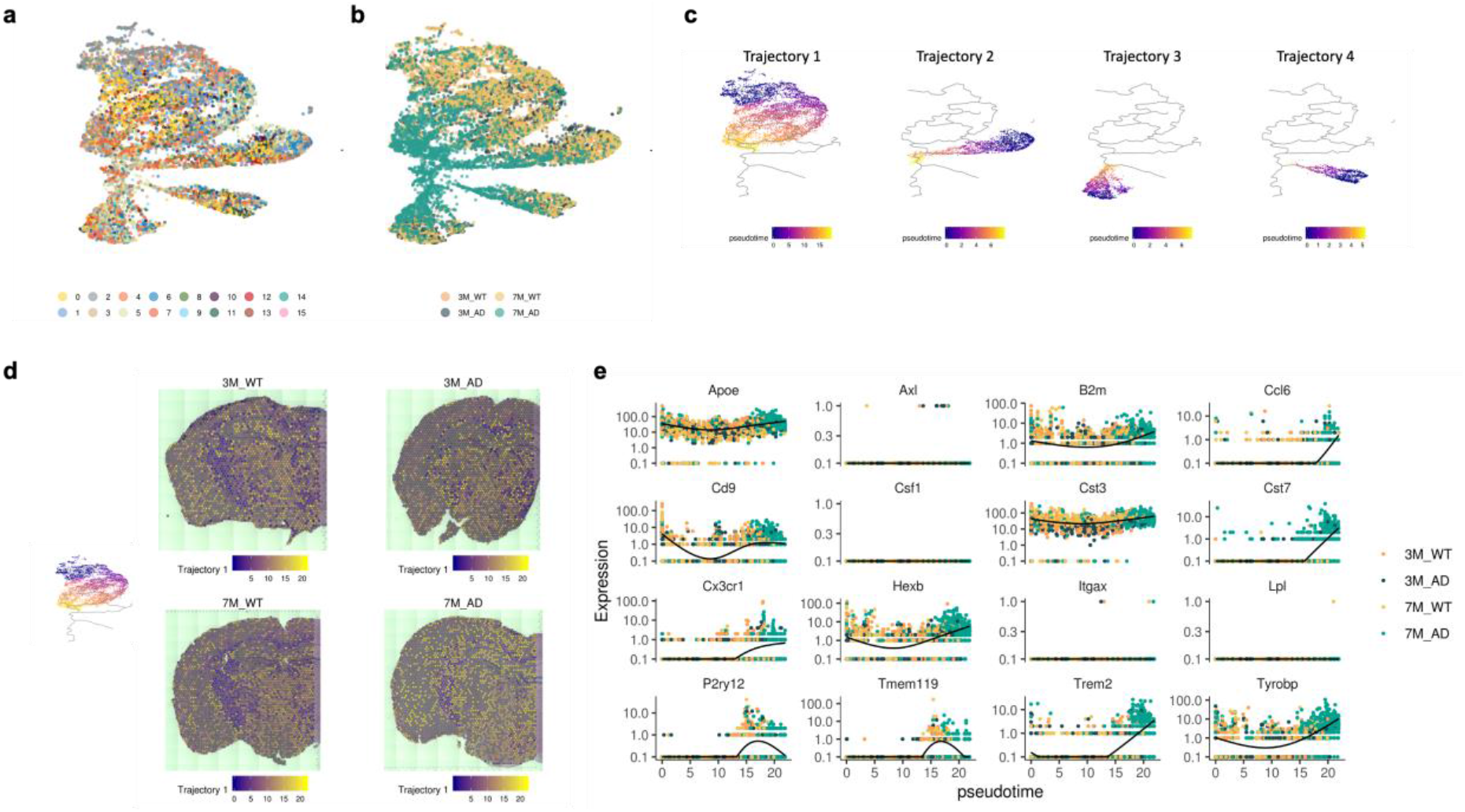
Trajectory inference of microglial signatures of spots. (a) UMAP represents microglial signature-based distribution. Notably, clusters determined by the transcriptome were mixed. (b) UMAP of microglial signatures colored according to mouse type is presented. (c) Trajectory inference analyzed with Monocle3 showed four different trajectories of spots of microglial signatures. (d) The first trajectory was represented with spatial maps and colored by pseudotime. Notably, the direction of pseudotime was determined by the predominant spots of the 7-month-old AD model in the center portion of UMAP. (e) The expression of microglial signature genes according to the pseudotime of trajectory 1 is presented. *Trem2, Ccl6,* and *Cst7* were increased at the late phase of pseudotime, while *Axl, Lpl*, and *Csf1* were negative until the late phase of pseudotime.

The activated status of trajectory 1, increased pseudotime, was found in both WT and AD models despite different spatial distributions (**Figure 6d**). The activated status of trajectory 1 was widely identified in most clusters of 7-month-old AD. However, 3-month-old AD brains showed an activated status of trajectory 1 in WM (**Figure 6d. Supplementary Fig. 13**). These results corresponded to previous results of early changes in WM. Notably, spots with an activated status of trajectory 1 were found in the hippocampus (cluster 4) and subdural area (cluster 8) in WT mice (**Supplementary Fig. 13**). In other words, trajectory 1 microglial activation was found in WT as well as AD, although AD showed spatially different patterns: WM at early stage (3 months) and the entire brain at late stage (7 months). Furthermore, activated status in WT identified in cluster 8 corresponded to relatively high DAM scores in our previous results (**Supplementary Fig. 10**). Gene expression according to the activated pattern of trajectory 1 is represented (**Figure 6e**). The activated status of trajectory 1 showed high *Ccl6, Cst7, Cx3cr1,* and *Trem2* and low expression of *Axl, Csf1,* and *Lpl*.

The other activated status of trajectory 2 was found in AD, while WT showed early pseudotime status of trajectory 2 (**Figure 7a**). The spatial distribution of trajectory 2 was also different from the spatial distribution of trajectory 1. In WT and 3-month-old AD model, spots with early pseudotime status were diffusely distributed in the thalamus and cerebral cortex (**Figure 7a**). The distribution was not changed in the thalamo-cortical regions of 7-month-old AD model. However, spots on the thalamus and cerebral cortex showed late pseudotime status of trajectory 2. Trajectory 2 showed high *Axl* expression regardless of the pseudotime, which was different from trajectory 1. Other markers, such as increased *Trem2* according to the pseudotime, were similar to the markers of trajectory 1 (**Figure 7b**).

**Figure 7.**
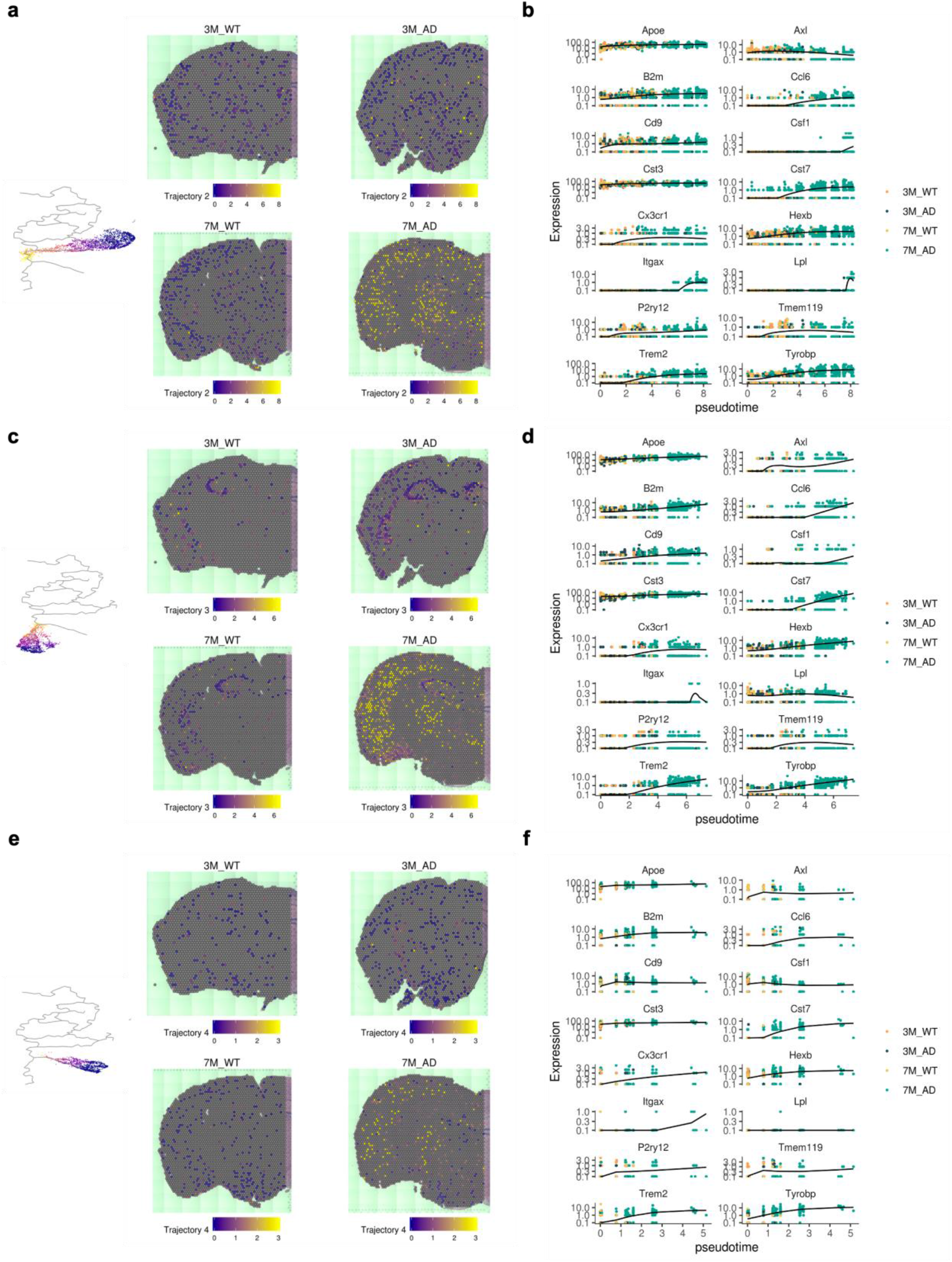
Trajectories of microglial signatures specific for AD. (a) Another trajectory, trajectory 2, was spatially mapped. The late phase of trajectory 2 was found only in the 7-month-old AD model, particularly thalamo-cortical regions. (b) The expression of microglial signature genes according to the pseudotime of trajectory 2 is represented. *Axl* was highly expressed in trajectory 2 regardless of pseudotime. (c) Trajectory 3 was spatially mapped. This trajectory commonly included spots of the hippocampus. The late pseudotime of trajectory 3 was found only in the 7-month-old AD model. (d) The expression of genes according to the pseudotime of trajectory 3. Notably, *Lpl* was highly expressed in trajectory 3 regardless of pseudotime. (e) Trajectory 4 showed sparse spots in the GM. The late pseudotime of this trajectory was found only in the 7-month-old AD. (f) Trajectory 4 was different from the other trajectories, as it had particularly high expression of *Csf1* regardless of pseudotime.

The activated status of trajectory 3 was also associated with AD, while WT and 3-month-old mice showed early status on trajectory 3 (**Figure 7c**). The spatial distribution was different from other trajectories. The spots included in trajectory 3 were identified in the hippocampus and amygdala. The activated status of trajectory 3 was found in the cortical cortex and thalamus of a 7-month-old AD model. In addition, spatial spots related to trajectory 3 pseudotime were identified in the striatum of early AD at 3 months (**Figure 7c**). Trajectory 3 was characterized by high expression of *Lpl* regardless of the pseudotime, which was different from other trajectories (**Figure 7d**). Of note, the spatial distribution of *Lpl* was increased in the hippocampus of WT according to in situ hybridization (ISH) data from the Allen Brain Atlas as well as spatial transcriptomic data (**Supplementary Fig. 14**).

Another trajectory 4 showed sparse distribution in the cortex and thalamus (**Figure 7e**). The late phase was found in the cerebral cortex and thalamus of a 7-month-old AD model. Spots of trajectory 4 were also identified in the WM of 3-month-old AD model, although it was early pseudotime status. Trajectory 4 was characterized by high expression of *Csf1* regardless of pseudotime (**Figure 7f**).

## DISCUSSION

We observed molecular features in the GM of AD models in the age-matched WT. The GM, including the cerebral cortex and hippocampus, showed a similar pattern of molecular changes. At 3 months, few molecular changes were observed, and there was a marked increase in reactive glial cell-related changes at 7 months. The dramatic activation of glial cells in the GM, an area where severe amyloid deposition occurs, demonstrated a close relationship with AD progression. Therefore, we concentrated on genes spatially associated with amyloid deposit patterns and investigated them by integrative analysis of the spatially resolved transcriptome and IF imaging. The top DEGs extracted by SPADE included genes related to glial activation, such as *Cst7* and *Ctsd*, while a few genes, such as *Prkcd, Rora*, and *Tcf712,* were not significantly upregulated genes in the AD model. These genes were associated with the regulation of ion transport. The spatial association suggested the possibility of functional interaction between amyloid deposits and membrane potential. Amyloid aggregation on the cell membrane results in lipid bilayer disruption and cell leakage, which are potential mechanisms of toxicity^20,21^. Of note, *Prkcd* was recently identified as a cerebrospinal fluid biomarker related to neurodegeneration induced by beta-amyloid^22^. In addition, we identified 20 genes that were correlated with amyloid deposit patterns and upregulated in GM at 7 months of AD. Among these genes, *Serpina3n* was different from the others and was involved mainly in microglial activation. Our results show that the expression level of *Serpina3n* gradually increased as AD pathology progressed dramatically in WM and GM. *Serpina3n* is commonly detected in astrocytes and activated oligodendrocytes and linked to increased amyloid accumulation, even though the mechanism is not clear23. According to a recent study, *Serpina3n* secreted by activated oligodendrocytes plays a role in promoting amyloid plaque deposition^24^. Furthermore, *Serpina3n* is a key marker of recently identified DAAs and is spatially adjacent to amyloid plaques^7^. These results confirmed that upregulated genes in AD spatially associated with amyloid plaques were consistent with previously identified markers functionally related to Aβ. Therefore, we suggest that changes in glial cells around amyloid plaques and changes in glial cells throughout the whole brain occur simultaneously.

An interesting biological finding in the current study is the initial changes in WM in AD models before distinct changes in GM. The major alteration of DEGs in GM was confirmed in 7-month-old AD mice, which had pronounced amyloidosis. In the GM at 7 months of AD, the expression levels of genes involved in reactive glial cells and lysosome pathways were upregulated compared to the expression levels of genes involved in reactive glial cells at 3 months (**Figure 2**). Of note, genes that were not significantly altered in GM but differently altered in WM were identified at 3 months. The early change before the upregulation of DEGs in GM could provide evidence of the role of WM changes independent of amyloid deposits in GM. This finding partly corresponds to the recent study with single nucleus RNA-seq analysis in 5XFAD mice, which identified a reactive oligodendrocyte population^24^. The study showed oligodendrocyte changes in 5XFAD mice, even in Trem2^−/−^ 5XFAD mice, even though they suggested that oligodendrocytes were affected by amyloid deposits. Additionally, a previous study with a spatial transcriptome in APP^NL-G-F^ mice suggested early alterations in a gene coexpression network related to myelin^25^. Our results suggested that WM changes functionally related to neuron projection and ensheathment of neurons were, at least, partly independent of amyloid plaques considering their early changes. Among the increased genes, *Mag* in WM was known as an inhibitor of axonal sprouting, which is critical for synapse formation (**Supplementary Fig. 6a**). This result provides evidence that myelin degradation may proceed first before pronounced amyloid accumulation. Furthermore, the WM of a 7-month-old AD model showed upregulation of genes related to reactive glial cells, as found in GM, such as *Trem2, Cst7,* and *Apoe* (**Figure 3c**). Thus, the results implied sequential molecular changes in WM: early changes independent of amyloid plaques followed by Trem2-dependent inflammatory signatures at the late phase.

In addition, AD-specific DAM and DAA signature analysis characterized the spatial distribution. Although the division of reactive glial cells into fixed categories is still controversial^26^, we used well-described activation markers to distinguish the reactive glial cells from homeostatic glial cells^5,7^. Our study showed that the signatures were increased exclusively in WM in the 3-month-old AD model. As pathology progressed, genes in disease-associated glial cells were upregulated in GM with more pronounced expression levels than in WM (**Figure 5**). This result is consistent with the finding of confirming the initially altered DEGs of WM in the 3-month AD model. Although the signatures are different from DAM, increased DAM in the WM at 3 months suggested Trem2-dependent microglial activation regardless of changes in GM. A recent report characterized white matter-associated microglia (WAMs) engaged in clearing myelin with single-cell RNA sequencing^13^. In an AD mouse model, WAMs displayed partial activation of the DAM gene signatures, allowing the identification of various aspects of DAM activity. WAMs, but no DAMs, appeared in 3-month-old APP^NL-G-F^ mice, which could reveal that myelin degeneration starts earlier than amyloid pathology^13^. Similar to microglia, DAAs showed the same activation patterns, initially from the WM to the thalamus and cortex area. In summary, the WM-to-GM transition provides important information about the pathological changes in AD progression, but further studies are needed to determine whether this transition occurs dependently and how glial cells communicate in each region.

Spatiotemporal changes in microglial signatures as key players in neuroinflammation were analyzed using a trajectory model. In this analysis, we selected spots containing genes related to DAM as well as homeostatic microglia and analyzed genes from all cells in the spot, including microglial signatures. Therefore, it is expected that not only changes in microglia but also the characteristics of the cells that change together were analyzed. As a result, the trajectory was divided into four distinct patterns (**Figure 6c**). The late phase of the distinctive trajectories reached a similar point associated with DAM genes. Trajectory 1 included both WT and AD despite different phase distributions of each group. However, trajectories 2 and 3 were found exclusively in the 7-month-old AD model. Trajectory 2 was associated with positive *Axl* expression, and trajectory 3 was associated with positive *Lpl* expression (**Figure 7**). Notably, *Lpl* expression was normally found in the hippocampus as an early phase of trajectory 3 (**Supplementary Fig. 14**). Although spatiotemporal patterns could not directly reflect microglia themselves, they suggested spatially distinctive activation patterns of glial cells. Even with the same DAM-related genes, there were genes that existed exclusively in AD pathology and were even present in WT. Our results suggest that hippocampus and subdural area in AD and even white matter area in WT have activated microglial signatures but were limited to trajectory 1 (**Supplementary Fig. 13**). These areas showed that *Axl* and *Lpl* were negative, indicating that they differed from AD-specific activated status. With a similar result, *Axl* expression was relatively high in only the 7-month-old AD model compared to *Itgax* and *Cst7*, which showed a significant increase even at 3 months (**Supplementary Fig. 9b**), indicating that *Axl* may be expressed at the late stage of AD pathology. The microglia that expressed the DAM signature genes appeared to have diverse activation patterns. Thus, these different microglia are distributed in specific brain regions, which is expected to provide information about the effects in different brain regions of AD. These results provide insight into which microglial changes we should pay attention to molecular targets in AD.

Overall, spatially resolved transcriptomic data from WT and AD mouse models of different ages showed spatiotemporally heterogeneous patterns of AD pathology. As cellular changes in a small region of the brain could not represent changes in the whole brain, these results provide resources for AD-related transcriptomic changes in gross-scale coverage with high-resolution spatial resolution.

## METHODS

### Mice

Three-month- and 7-month-old male 5XFAD mice (Tg6799; on a C57/BL6-SJL background) containing five FAD mutations in human APP (the Swedish mutation, K670N/M671L; the Florida mutation, I716V; and the London mutation, V717I) and PS1 (M146L/L286V) and wild-type mice were used for spatially resolved transcriptomic data and quantitative PCR. Male 5XFAD mice aged 2, 4, 7.5 and 12 months were used for IF imaging of tissue sections. All experimental protocols and animal usage were approved by the Institutional Animal Care and Use Committee at Seoul National University.

### Immunofluorescence imaging for tissue sections

Paraffin-embedded brain tissues were sectioned at 4-μm thickness. Deparaffinization was achieved with xylenes and decreasing concentrations of ethanol. Tissue sections were subjected to antigen retrieval using citrate buffer, pH 6.0, at boiling temperature for 10 min. Following rinsing with TBS, sections were incubated in blocking buffer containing TBS with 0.5% BSA for 1 h at room temperature. Slides were then incubated with primary antibody in blocking buffer overnight at 4 °C. The next day, slides were washed with TBS and then stained with Alexa Fluor secondary antibodies (Thermo Fisher Scientific). Sections were rinsed again and stained with DAPI (1:100; Invitrogen) before being cover-slipped with mounting medium. The primary antibodies used were rabbit Iba1 (1:200; Abcam), rabbit LAMP1 (1:50; Abcam), mouse β-Amyloid (6E10) (1:100; BioLegend), rabbit β-Amyloid (D54D2) XP (1:100; Cell Signaling Technology), mouse GFAP (1:200; Cell Signaling Technology), and rabbit dMBP (1:100; Sigma Aldrich).

### Thioflavin S staining

The paraffin-embedded sections were deparaffinized in xylene and rehydrated in ethanol solution. The hydrated brain sections were incubated in 1% thioflavin S solution for 5 minutes and washed with 70% ethanol and distilled water. For the images of stained slides, LEICA confocal microscopy SP8 was used.

### Microdissection of brain tissue and quantitative PCR (qPCR)

Brains were collected and microdissected to obtain samples of the WM and the cortex (free of meninges and choroid plexus). RNA extraction was performed using TRIzol (Thermo Fisher) according to the manufacturers’ protocols. Reverse transcription of RNA was performed using a thermal cycler (Bio-Rad, T100). cDNA samples were diluted and mixed with SYBR green master mix (Takara) before loading as technical triplicates for qPCR on an Applied Biosystems 7500. Primers are specified in **Supplementary Table 4**.

### Spatially resolved transcriptomic data generation

Prepared brain hemispheres were cryosectioned to thin (10 μm) coronal sections and processed the same day. First, mouse brain sections were sectioned and mounted onto slides on Visium Spatial Tissue Optimization slides (10x Genomics). The tissue permeabilization time was determined by the manufacturer’s protocols (VisiumSpatialTissueOptimization, https://support.10xgenomics.com/spatial-gene-expression/tissue-optimization). Accordingly, tissue was permeabilized for 12 min for Visium Spatial Gene Expression analysis. Before library preparation, tissue sections were methanol-fixed, hematoxylin and eosin (H&E) -stained and imaged on a TissueFAXS PLUS (TissueGenostics). The slides were merged into a picture of the whole brain using TissueFAXS imaging software. Sections were then permeabilized and processed to obtain cDNA libraries. cDNA libraries were prepared according to the manufacturer’s protocol (VisiumSpatialGeneExpression, https://support.10xgenomics.com/spatial-gene-expression/library-prep). To verify the size of PCR-enriched fragments, we checked the template size distribution by running on an Agilent Technologies 2100 Bioanalyzer. The libraries were sequenced using HiSeqXten (Illumina) with a read length of 28 bp for read 1 (Spatial Barcode and UMI), 10 bp index read (i7 index), 10 bp index read (i5 index), and 90 bp for read 2 (RNA read).

Raw FASTQ data and H&E images were processed by the Space Ranger v1.1.0 (10X Genomics) pipeline for the gene expression analysis of Visium spatial gene expression library data. Illumina base call files from the Illumina sequencing instrument were converted to FASTQ format for each sample using the ‘mkfastq’ command. Visium spatial expression libraries were analyzed with the ‘count’ command. Image alignment to predefined spots was performed by the fiducial alignment grid of the tissue image to determine the orientation and position of the input image. Sequencing reads were aligned to the mm10 reference genome (mm10-2020-A) using STAR (v2.5.1b) aligner. Gene expression profiling in each spot was performed with unique molecular identifier (UMI) and 10X barcode information.

### Spatial Transcriptomics Data Clustering

The spots with gene expression data were analyzed with the Seurat package (ver 3.1.2) ^27^. Gene counts were normalized using ‘LogNormalize’ methods in Seurat v.3. The top highly variable genes (n= 2,000) were then identified using the ‘vst’ method in Seurat. The number of RNA counts for each spot and the frequency of mitochondrial gene counts were regressed out in the scaling process. Four spatial transcriptomic datasets were merged and rescaled. Principal component analysis was performed using the top highly variable genes. For visualization, dimension reduction was performed using UMAP on the top 30 principal components. Graph-based clustering based on the Louvain community detection algorithm was performed. The resolution was set to 0.5. Markers for each cluster were identified by Wilcoxon rank sum tests for a given cluster vs. other clusters implemented in Seurat as a ‘FindAllMarkers’ function. The anatomical location of each cluster was visually identified by comparison with the Allen Mouse Brain Reference Atlas (https://mouse.brain-map.org/static/atlas).

### Differential gene expression analysis on clustering of spots

MAST28 in the Seurat function was used to perform differential gene expression analysis. First, we identified differentially expressed genes between two specific clusters (for this study, cluster 7 vs. cluster 1, two clusters of thalamus). A gene was considered significant with false discovery rate (FDR) -adjusted P < 0.05 and log-fold change (logFC) > 0.3. Volcano plots were drawn by EnhancedVolcano function in R. In addition, differentially expressed genes were extracted from the comparison of AD and WT mice at 3 months and 7 months. The analysis was also performed using the MAST function after selecting a subset of specific clusters. The cutoff of significantly different genes was FDR-adjusted p < 0.05 and log FC > 0.25. Gene ontology and KEGG pathway analyses were performed with clusterProfiler^29^, which supports statistical analysis and visualization of functional profiles for genes and gene clusters.

### Microglial and astrocytes signature scores

Gene sets of microglia and DAM were selected for the matrices of spatial transcriptomic data to calculate the signature score. The score was calculated with the AddModuleScore function with default parameters in Seurat. DAMscore was visualized by the SpatialFeaturePlot function for identifying spatial distribution patterns. In addition, gene sets of DAAs were selected to calculate the signature score. DAA scores were also visualized by the SpatialFeaturePlot function.

### Spatial coregistration of the immunofluorescence image

The immunofluorescence image of amyloid plaque (6E10; mouse) obtained from a 7-month-old AD model was spatially coregistered with the H&E image obtained before spatial transcriptomic data acquisition, i.e., Visium library generation. Notably, the immunofluorescence image of amyloid plaque was the adjacent slide of H&E image for the spatial trarnscriptomic data generation. The immunofluorescence image was resized to have a similar size of H&E images. Cropping and rotation were performed to approximately overlap both images. This process was performed by the scikit-image package (version 0.15.0). For image registration, both images were preprocessed by: 1) changing to grayscale, 2) masking with a pixelwise threshold to include the mouse brain, and 3) smoothing using a Gaussian filter (sigma value of 5). The transform function for the coregistration was estimated using the Dipy package (version 1.0.0). The image was linearly transformed by rigid and affine transformation. For the final coregistration, nonlinear warping was performed using SymmetricDiffeomorphicRegistration based on the Symmetric Normalization (SyN) algorithm30. After the estimation of transformation function, the immunofluorescence image was transformed for further analysis integrating image patterns and spatial transcriptomic data.

### Integrative analysis of the immunofluorescence image and spatial transcriptomic data

Molecular features associated with tissue image patterns were extracted by the SPADE tool^18^. The coregistered amyloid plaque immunofluorescence image was used for the input of SPADE. The CNN-derived image features (VGG-16 model) were extracted by 32 × 32 sized patches centered at the spots. For dimension reduction, principal components of image features were used. In this study, the first principal component represented the accumulation of amyloid plaques; thus, genes correlated with the first principal component of image patterns were identified. The top 100 genes according to the regression coefficient (logRC) were selected. Gene ontology analysis was performed with clusterProfiler.

### Spatiotemporal trajectory using pseudotime analysis

A gene set with microglial signatures was selected to generate a matrix for spatially resolved gene expression data: *Hexb, Cst3, Cx3cr1, Ctsd, Csf1r, Ctss, Sparc, Tmsb4x, P2ry12, C1qa, C1qb, Tmem119, Tyrobp, Ctsb, Apoe, B2m, Fth1, Lyz2, Trem2, Axl, Cst7, Ctsl, Lpl, Cd9, Csf1, Ccl6, Itgax,* and *Timp2*5,6,31. Normalization and PCA were performed using the ‘preprocess_cds’ function from Monocle version 3. The trajectory graph based on the microglial signature gene set was learned by the ‘learn_graph’ function from Monocle. The UMAP plots according to the trajectory were drawn with colors of the clusters using all genes analyzed with Seurat as well as mice. Subsequently, the spots were semiautomatically ordered according to the progression of microglia. The trajectory was automatically learned; however, the direction of order was determined by UMAP plot with mice (AD vs WT). As the spots of 7-month-old AD mice were located at a specific portion, the center of the UMAP, the spots enriched in 7-month-old AD mice were regarded as ‘late-phase’ pseudotime.

Spatial mapping of pseudotime for each trajectory was performed by selecting the subset of spots included in the selected trajectory. Colors with pseudotime were mapped on the spots of specific locations and mapped using SpatialFeaturePlot from Seurat.

### Statistics

Differentially expressed genes were identified by the aforementioned methods. All P values reported in this study were two-sized and adjusted by false discovery rate. The statistical method incorporated in the specific software was used with a default parameter unless otherwise indicated. Plots in R were created either with the ggplot2 R package or Seurat modified by custom codes for data visualization.

## DECLARATIONS

### Acknowledgement

The work was supported by grants from the National Research Foundation of Korea (NRF-2017M3C7A1048079, NRF-2019R1F1A1061412, NRF-2019K1A3A1A14065446, NRF-2020M3A9B6038086, NRF-2020R1A2C2101069).

### Competing interests

The authors declare no competing financial interests.

### Author contributions

H.C., Y.C., D.S.L designed and supervised the study. H.C. developed the custom code for analysis. E.J.L. performed spatially resolved transcriptomic data generation. H.C. and S.B. processed and analyzed the sequencing data. E.J.L., J.S.S., K.H., performed immunofluorescence staining and qRT-PCR experiments. H.C., E.J.L., Y.C., D.S.L. wrote the paper. All authors read and approved the final manuscript.

## Notes

### Competing Interest Statement

The authors have declared no competing interest.

