## Supplementary Figures for "Spatiotemporal characterization of glial cell activation in an Alzheimer’s disease model by spatially resolved transcriptome"

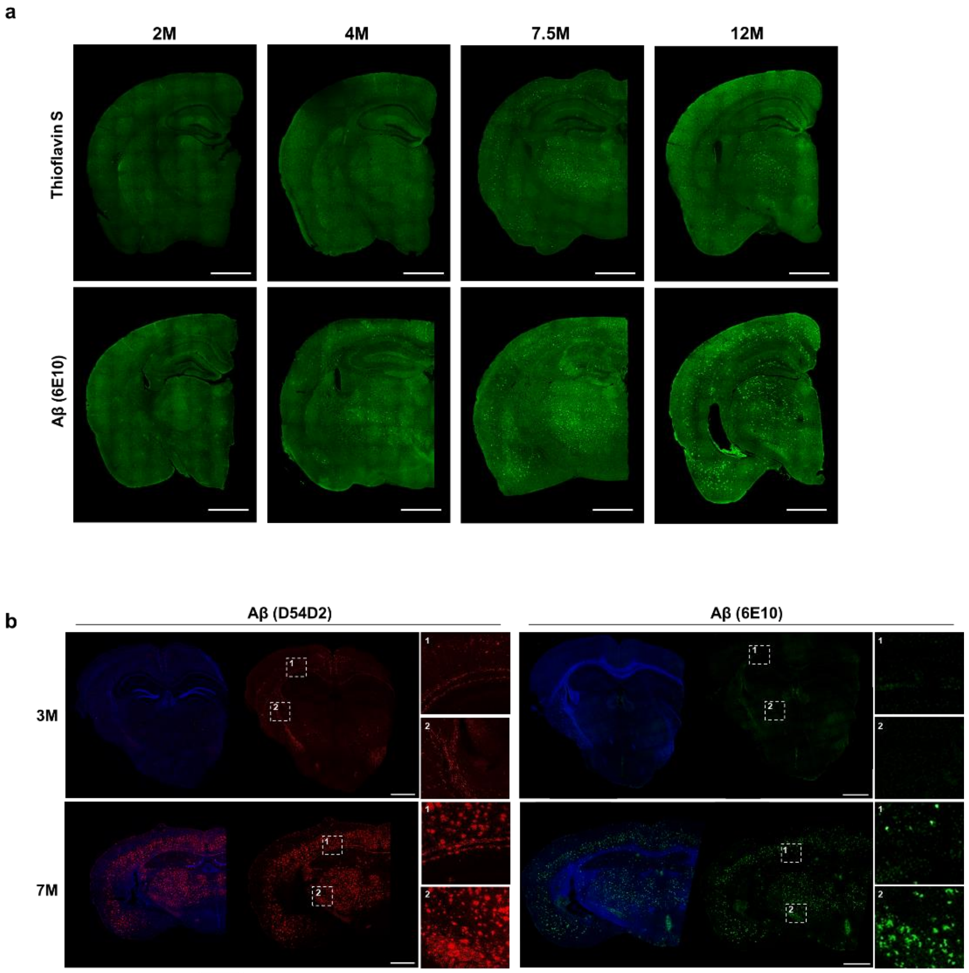


**Supplementary figure 1. Age-dependent accumulation of β-amyloid in AD model brains**

(a) Immunofluorescence imaging on whole brain tissue slides from 2, 4, 7.5 and 12-months-old AD model is shown (n=3). Coronal brain sections were stained with Thioflavin S (*above*) and 6E10 antibody showing Aβ accumulation (*below*). (b) Immunofluorescence imaging for the brain tissues, 3-and 7-month-old 5XFAD, is shown. These images were acquired from the same brain from which spatially resolved transcriptome data were acquired. Aβ was stained with D54D2 antibody (red) and 6E10 antibody (green), and Aβ accumulation in the GM of the 3-month-old AD model was observed. In the inset, 1 indicates corpus callosum and cortex, 2 indicates internal capsule and thalamus. The images were acquired using tile-scaling LEICA confocal imaging software. Scale bars, 1 mm. (2M: 2-month-old; 4M: 4-month-old; 7.5M: 7.5-month-old; 12M: 12-month-old)

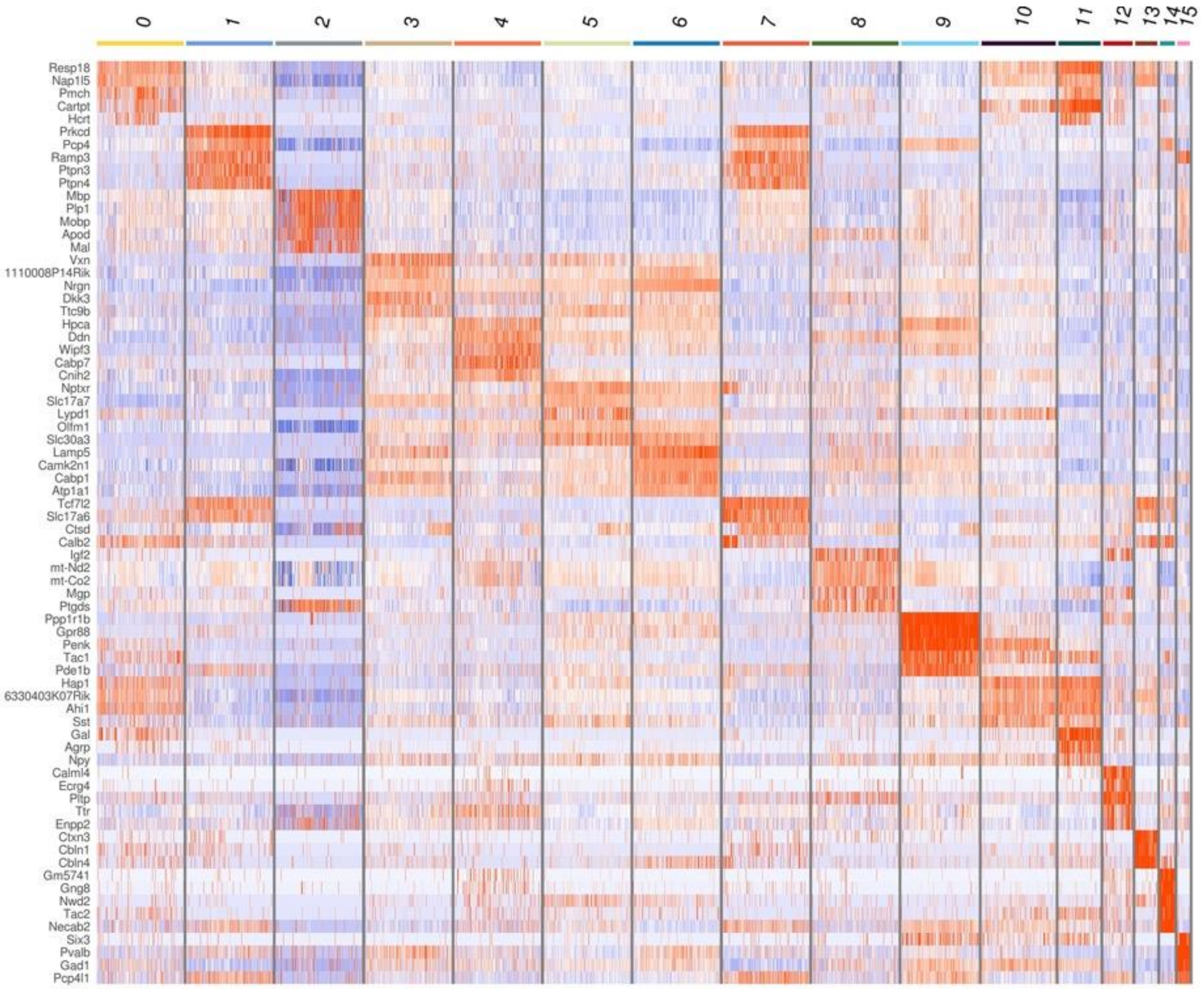


**Supplementary figure 2. Marker genes of spots of spatial transcriptome.**

Spots of spatial transcriptome of WT and AD models were clustered according to gene expression. (WT: wild type; AD: Alzheimer’s disease)

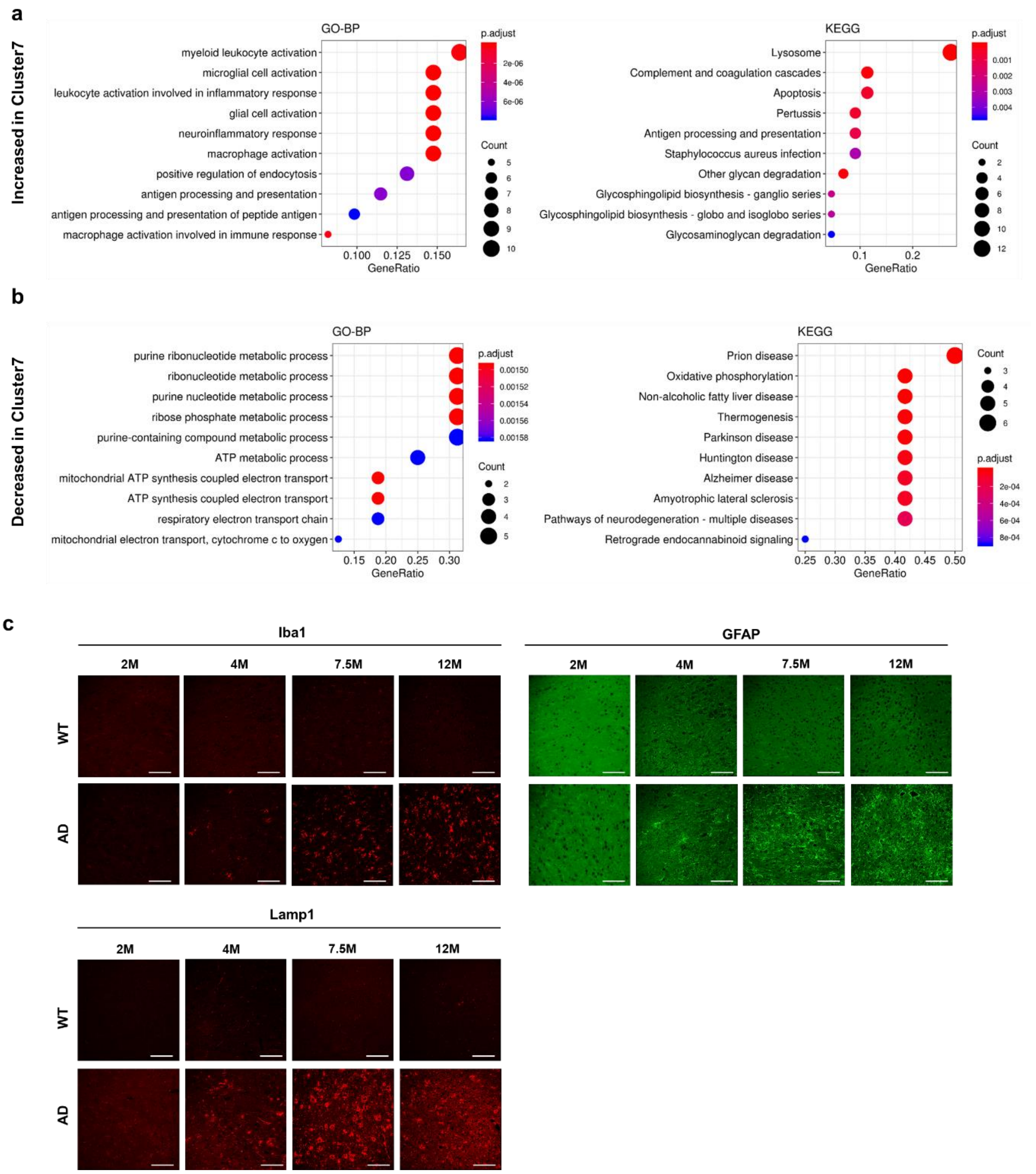


**Supplementary figure 3. The gene ontology of cluster 7 specifically found in AD model at 7 months.** (a) GO and KEGG pathway analyses of upregulated genes of cluster 7 compared with cluster 1 which represented thalamus of WT and 3-month-old AD model. (b) GO and KEGG pathway analyses of downregulated genes of cluster 7 compared with cluster 1. (c) Immunostaining of microglia/macrophages (Iba1, red), astrocytes (GFAP, green), and lysosomal function (Lamp1, red) were observed in thalamus at the indicated ages (n=3). Scale bars, 50 μm.

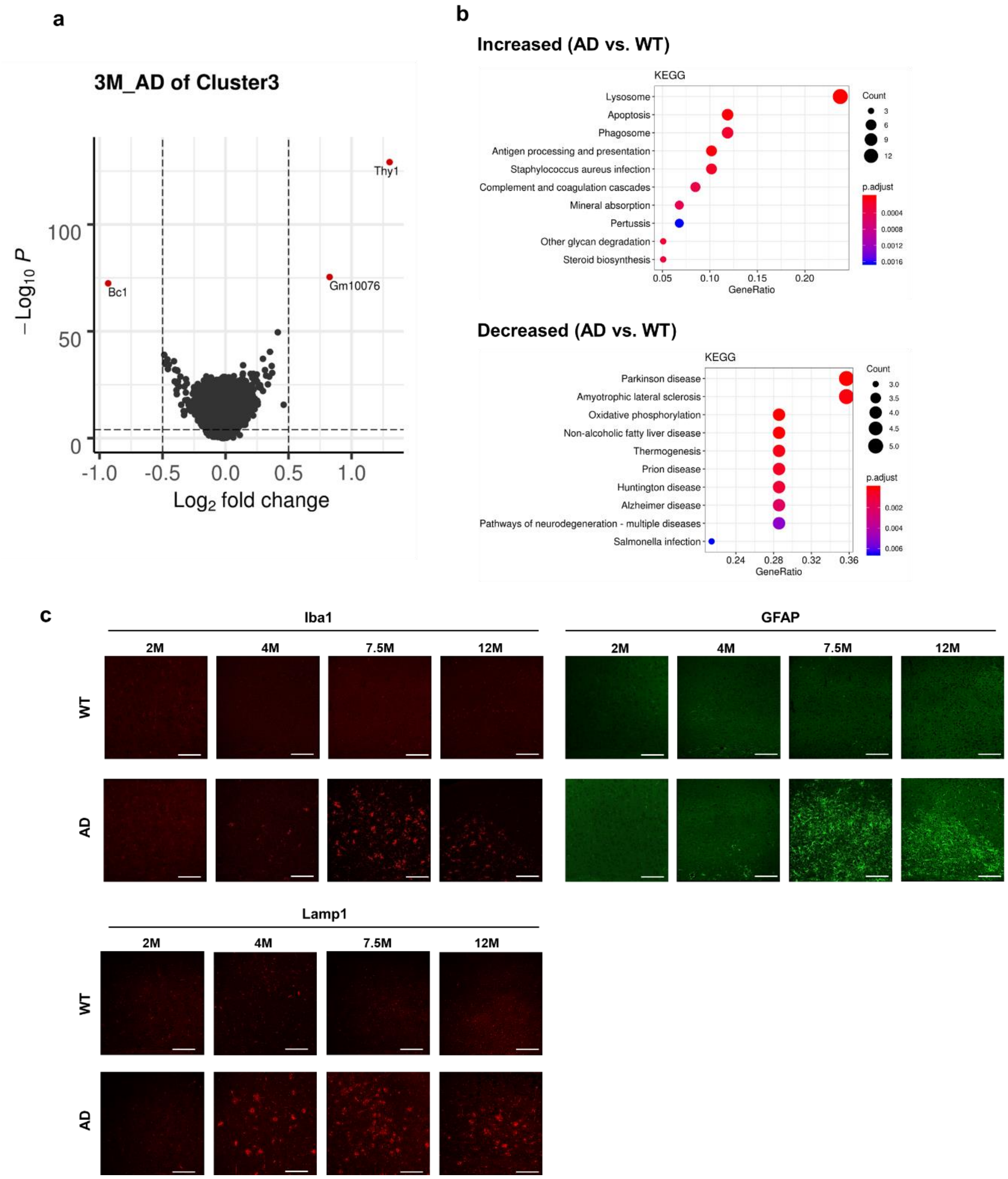


**Supplementary figure 4. Differentially expressed genes in the cerebral cortex of AD model, cluster 3.** (a) Differentially expressed genes in 3-month-old AD model compared with WT. (b) KEGG pathways of differentially expressed genes in 7-month-old AD model. (c) Immunostaining of microglia/macrophages (Iba1, red), astrocytes (GFAP, green), and lysosomal function (Lamp1, red) were observed in cortex at the indicated ages (n=3). Scale bars, 50 μm.

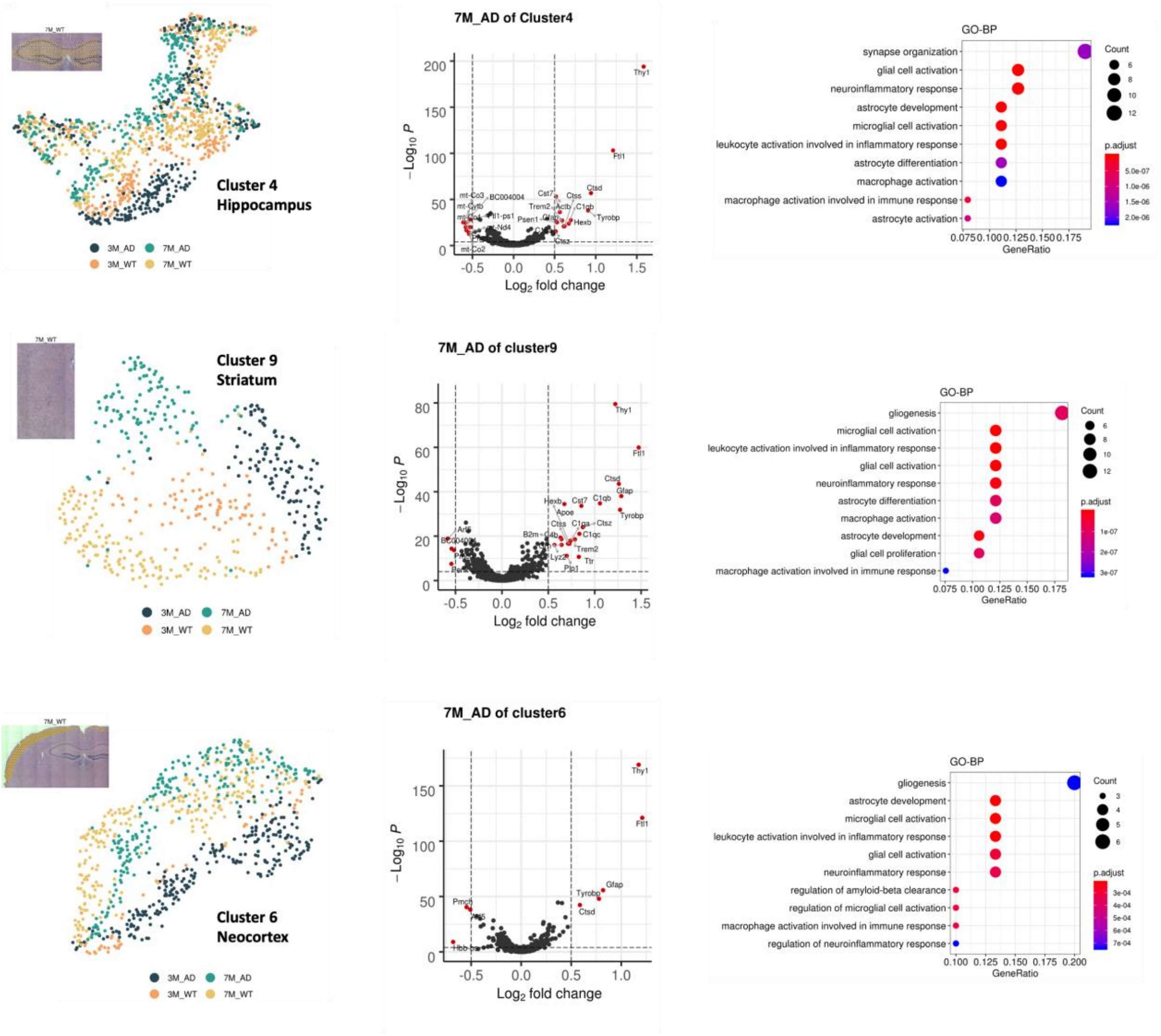


**Supplementary figure 5. Differentially expressed genes and GO of the AD model according to clusters.**

Differentially expressed genes and GO terms of upregulated genes in 7-month-old AD model compared with WT. The analyses were performed in cluster 4 (hippocampus), cluster 9 (striatum), and cluster 6 (outer layer of cerebral cortex).

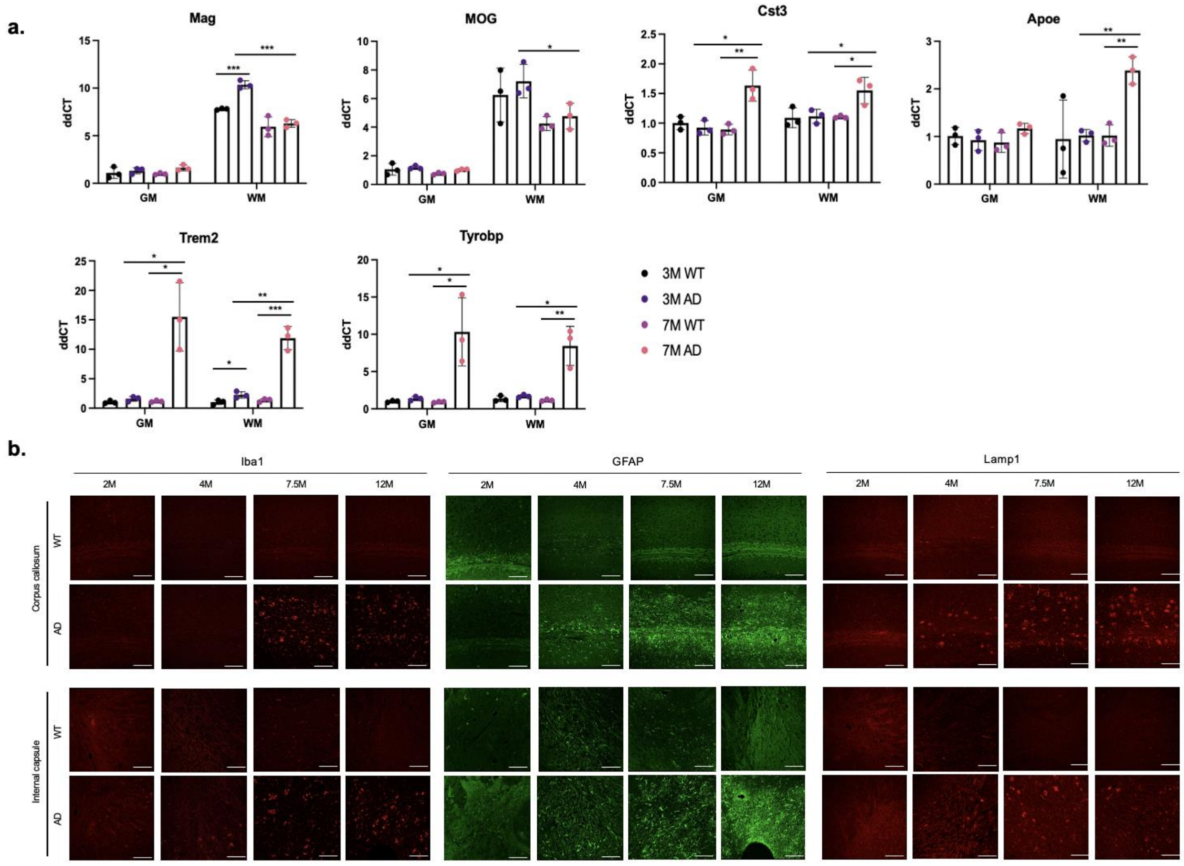


**Supplementary figure 6. Immunofluorescence images of white matter**

(a) qPCR analysis of *Mag, MOG, Cst3, Apoe, Trem2, and Tyrobp* in the different groups of mice. Each symbol represents an individual mouse. *p<0.05, **p<0.01, ***p<0.001 (one-way ANOVA). Scale bars, 50 μm. (b) Immunostaining of microglia/macrophages (Iba1, red), astrocytes (GFAP, green), and lysosomal function (Lamp1, red) were observed at the indicated ages (n=3). Notably, images showed increased reactive astrocytes, microglia, and lysosomal function in WM, internal capsule and corpus callosum, at 4-month AD model. Scale bars, 100 μm.

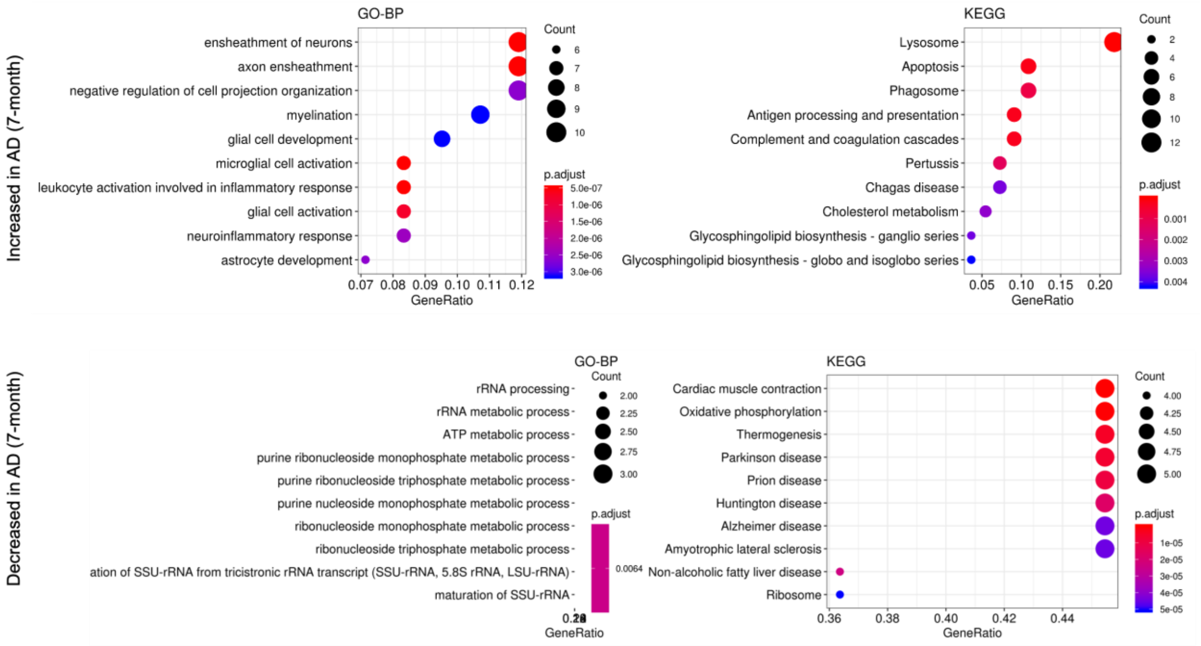


**Supplementary figure 7. Functional term of differentially expressed genes of cluster 2 (white matter) in 7-month-old AD model.**

GO terms and KEGG pathways of upregulated genes in 7-month-old AD model were represented (*above*). In addition, those of downregulated genes in 7-month-old AD model were also represented (*below*).

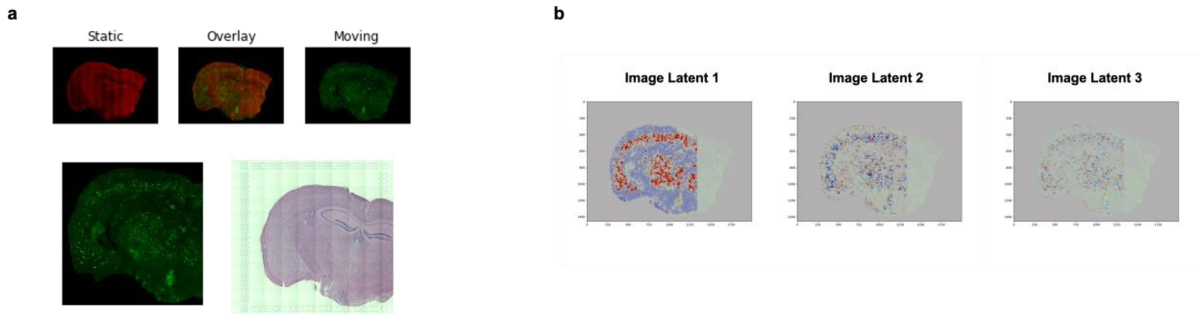


**Supplementary figure 8. Registration of the IF image to H&E and image latent features.**

(a) The IF image of 6E10 antibody represented Aβ accumulation acquired from 7-month-old AD model, a same animal with spatially resolved transcriptome. The IF image was coregistered with H&E image obtained for spatial transcriptomic data. (b) The spatial distribution of 2nd and 3rd image latents, derived from SPADE algorithm, was presented. These distribution patterns were visually different from Aβ accumulation.

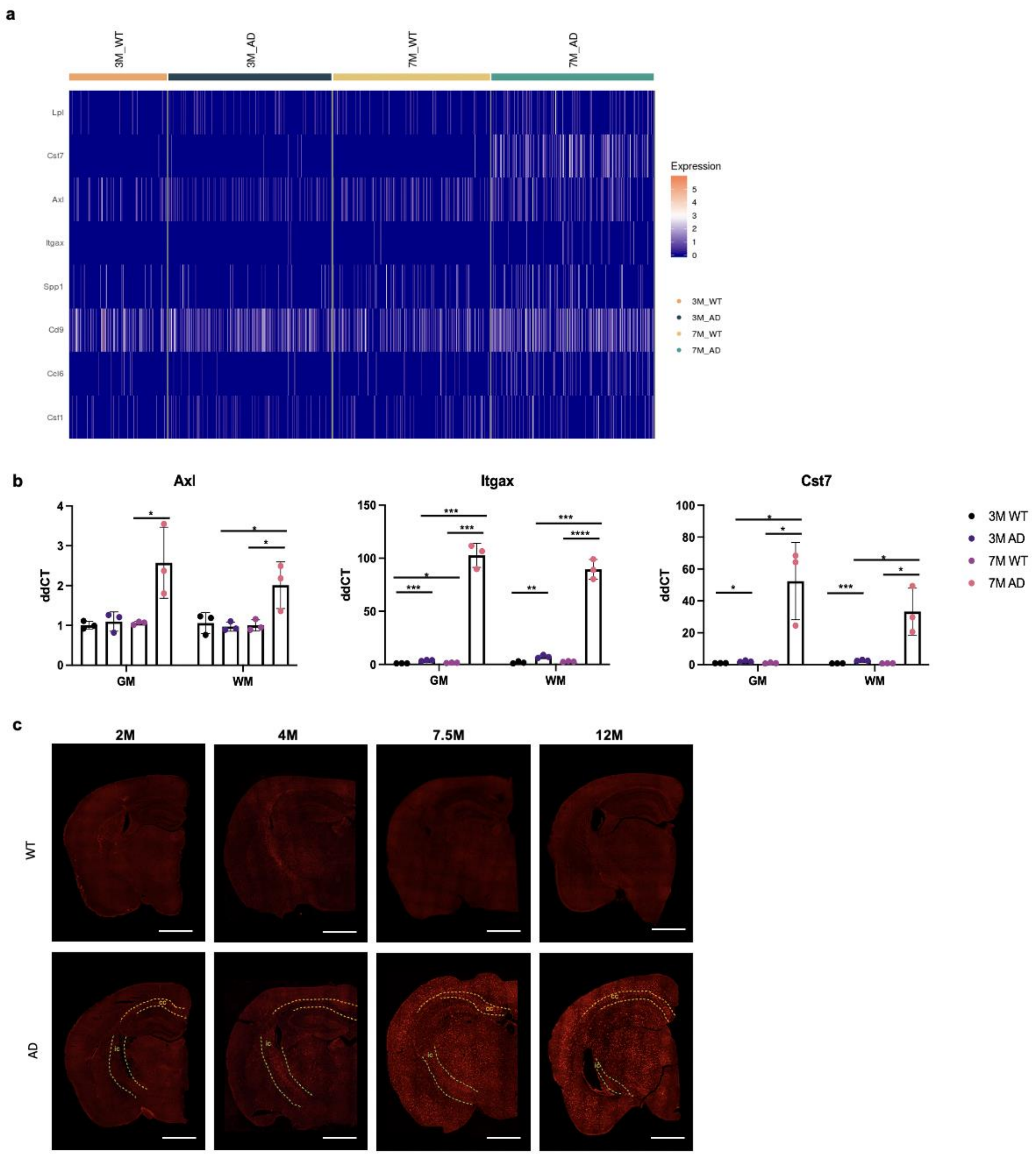


**Supplementary figure 9. Disease-associated microglia (DAM) signatures and temporal IF images for microglia**

(a) A heatmap for DAM signature genes according to mice. (b) qPCR analysis of *Axl, Itgax, and Cst7* in the different groups of mice. Each symbol represents an individual mouse. *p<0.05, **p<0.01, ***p<0.001 (one-way ANOVA). Scale bars, 1 mm. (c) IF with anti-Iba1 revealed increased microglia according to aging in the AD model (n=3). Notably, fluorescence signal was identified in internal capsule and corpus callosum in 4-month AD model, and the activity was clearly increased in cortex and thalamus at 7.5 months.

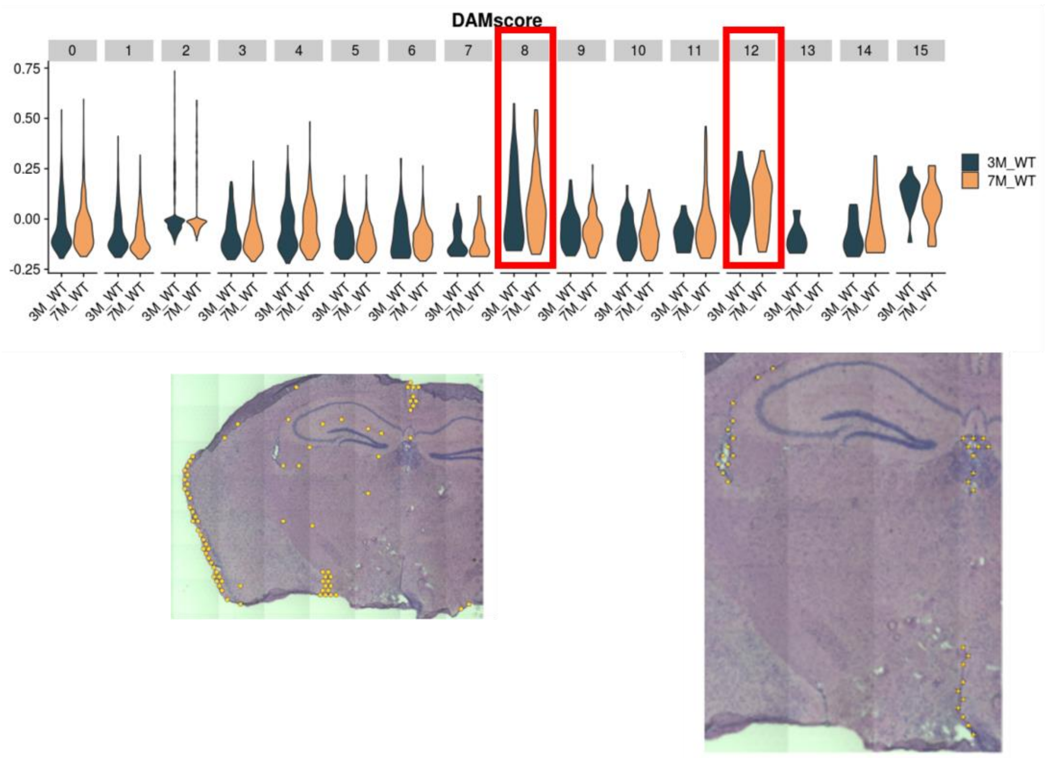


**Supplementary figure 10. DAM scores in WT mice.**

Two clusters of WT-mice showed relatively high DAM score. These brain regions subdural area (cluster 8) and periventricular area (cluster 12).

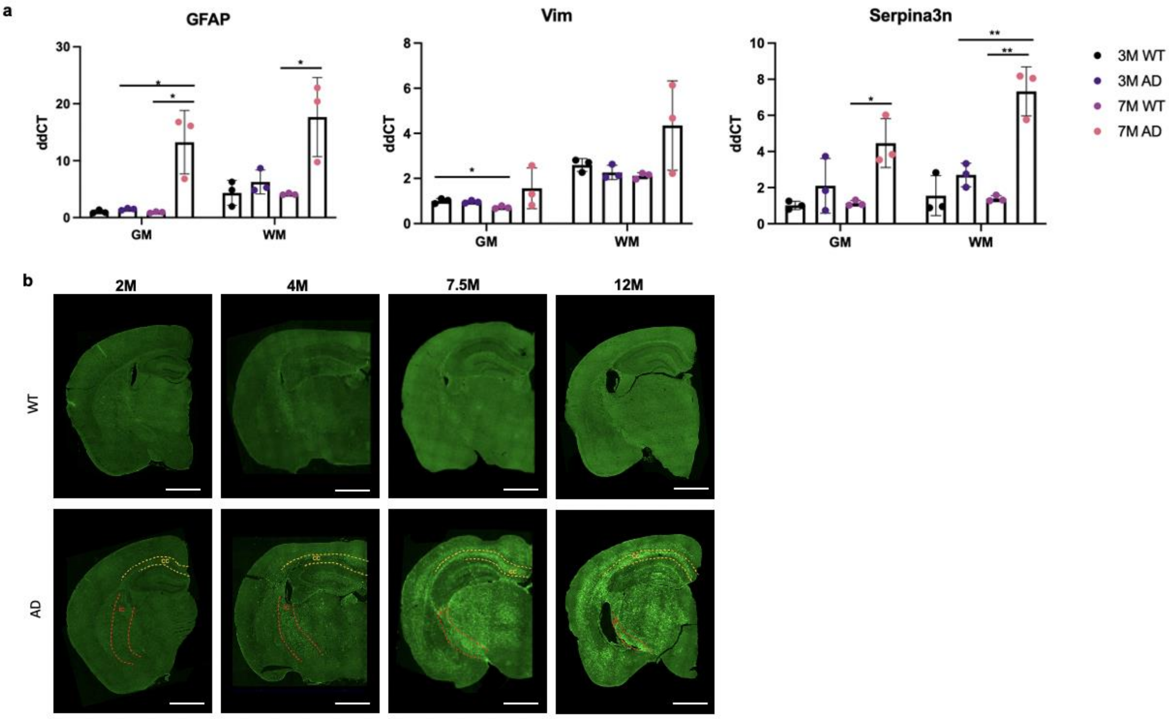


**Supplementary figure 11. IF image for astrocytes** (a) qPCR analysis of *Gfap, Vim, and Serpina3n* in the different groups of mice. Each symbol represents an individual mouse. *p<0.05, **p<0.01, ***p<0.001 (one-way ANOVA). Scale bars, 1 mm. (b) IF with anti-GFAP revealed increased the activation of astrocytes according to aging in the AD model (n=3). Notably, fluorescence signal was identified in internal capsule and corpus callosum in 4-month AD model, and the activity was clearly increased in cortex and thalamus at 7.5 months.

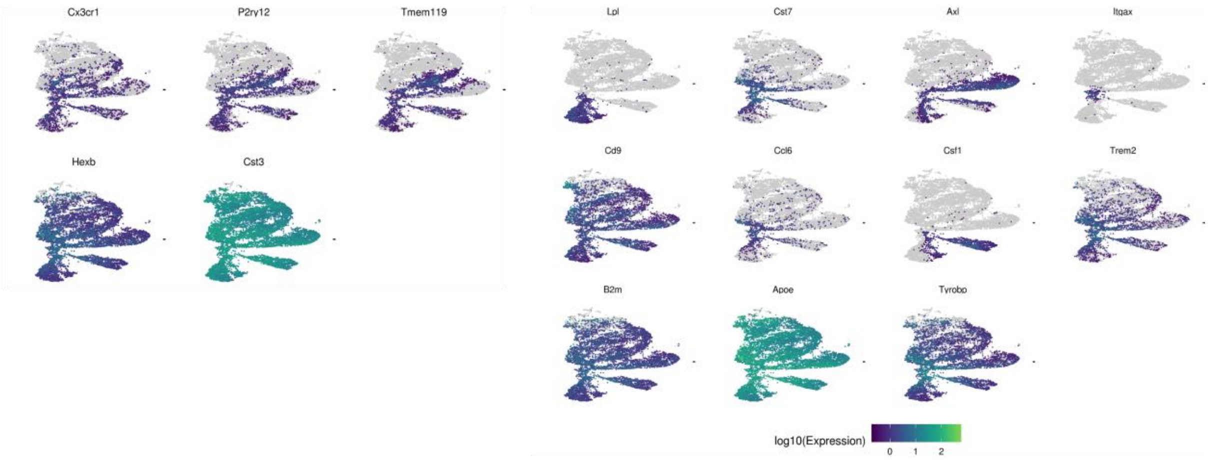


**Supplementary figure 12. The expression of microglial genes.**

The expression of key genes of microglia was represented by a color map with log10(Expression) with UMAP plots drawn by microglial gene sets.

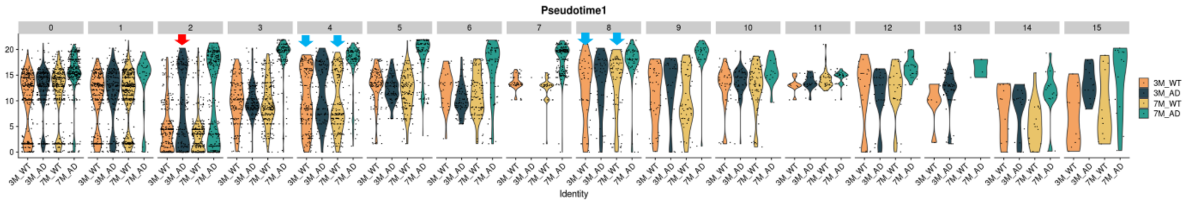


**Supplementary figure 13. The pseudotime of 'trajectory 1' of microglia according to mice.** The pseudotime of trajectory 1 represented microglial activation in various clusters of the 7-month-old AD model. Notably, the relatively high pseudotime was found in cluster 2 (White matter) of 3-month-old AD model (Red Arrow). In addition, spots with relatively high pseudotime was found in WT. These spots were included in cluster 4 (hippocampus) and cluster 8 (subdural area) (Blue arrows).

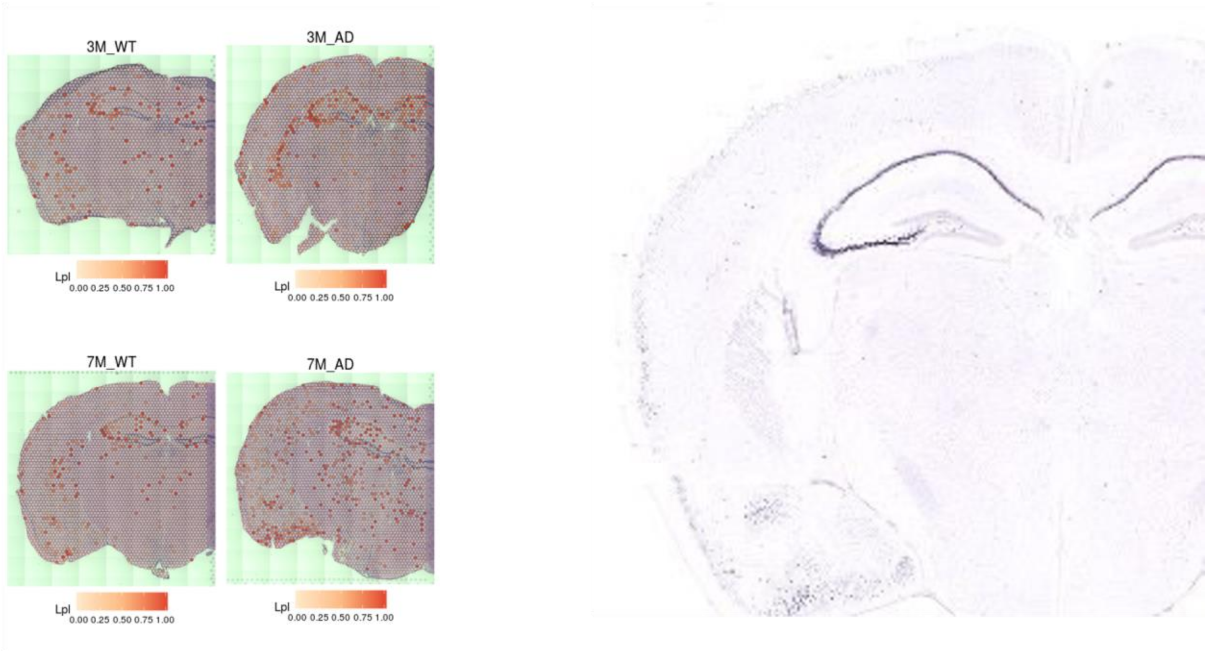


**Supplementary figure 14. Spatial distribution map of *Lpl*.**

The trajectory 3 was associated with AD-specific high pseudotime and characterized by high *Lpl*. *Lpl* expression was found in the hippocampus of WT as well as AD (*left*). This spatial pattern of *Lpl* expression was also identified by in situ hybridization data (ISH) from the Allen Brain Atlas (*right*).
